# Revealing year-round activity of cave-dwelling insectivorous bats with a sonotype classifier in data-deficient areas

**DOI:** 10.1101/2025.02.22.639615

**Authors:** Morgane Labadie, Serge Morand, Alexandre Caron, Helene Marie De Nys, Fabien Roch Niama, Franel Nguilili, N’Kaya Tobi, Mathieu Bourgarel, Yves Bas, Charlotte Roemer

**Affiliations:** CIRAD, UMR ASTRE, F-34398 Montpellier, France; ASTRE, CIRAD, INRAE, Univ Montpellier, Montpellier, France; IRL HealthDEEP CNRS – Kasetsart University – Mahidol University, Bangkok, Thailand; Faculty of Veterinary Technology, Kasetsart University, Bangkok, Thailand; Department of Social and Environmental medicine, Faculty of Tropical Medicine, Mahidol University, Bangkok, Thailand; Faculdade de Veterinaria, Universidade Eduardo Mondlane, Maputo, Mozambique; CIRAD, UMR ASTRE, Harare, Zimbabwe; Laboratoire National de Santé Publique, Brazzaville, République du Congo; Direction Générale de l’Élevage (Service vétérinaire), Ministère de l’Agriculture, de l’élevage et de la pêche, Brazzaville, République du Congo; Centre d’Ecologie et des Sciences de la Conservation (CESCO), Muséum National d’Histoire Naturelle, Centre National de la Recherche Scientifique, Sorbonne Université, Paris, France; CEFE, Univ Montpellier, CNRS, EPHE, IRD, Montpellier, France

**Keywords:** PAM, Tadarida, activity pattern, abundance, bats phenology, Congo

## Abstract

**Context:** Bats are the only mammals who have conquered the skies and the leading mammals in terms of total biomass with the exception of humans and domestic mammals. However, the ecology of many bat species is still unknown, especially in poorly sampled regions such as Central Africa.

**Aims:** We present the first application of an acoustic method for studying the nocturnal activity patterns and annual phenology of bat activity of cave bat communities in regions where no bat acoustic classifier exists.

**Methods:** In two caves (Mont Belo and Boundou) in the Republic of Congo, which are home to several species of insectivorous bats throughout the year, we set up a passive acoustic monitoring (PAM) over a period of 19 months. To identify the massive recordings collected, we used a sonotype classifier. To assign a species name or species group to the different sonotypes identified by the classifier, we collected local reference calls by capturing bats at the cave entrances. This enabled us to identify two bat groups and three species despite the absence of local call libraries. The capture sessions also allowed us to collect information about the reproductive condition of the individuals, which we interpreted in relation to the annual phenology of bat activity. To quantify bat activity, we defined the activity of each acoustic group as the number of bat passes, where a bat pass corresponds to the detection of one or several echolocation calls within a five-second interval. This standardized measure allowed us to quantify nightly and seasonal variations in activity.

**Key results:** The nightly activity displayed strong peaks in the evening and the morning, except during the long rainy season, when a tri- or quadri-modal activity was detected. This more evenly distributed activity throughout the night is related to the period of juvenile rearing (gestation, parturition and lactation). For all acoustic groups, we also observed a higher activity during the rainy seasons and the short dry season, compared to the long dry season. This high activity is linked to the lactation period, the independence of young bats and potentially the mating period. Finally, the reproductive activity of insectivorous bat species is synchronized with the period of high resource availability.

**Conclusion:** This study represents the first long-term acoustic monitoring of cave bats in the Republic of Congo. This was allowed by the use of a classifier built without any pre-existing regional call data, highlighting a practical pathway for studying bat activity in other data-deficient regions.

**Implications:** Information about the biological status of individuals captured at the monitored caves in association with the cave annual activity phenology produced with this method are key to progress in the understanding of the ecology and the conservation issues of cave bat communities, which are particularly threatened globally.

## Introduction

Human activities are currently triggering a massive loss of biodiversity, with rates of species decline or extinction unprecedented for several millennia (Barnosky et al., 2011; Ceballos et al., 2015; IPBES, 2019). However, the scale of this crisis is probably still concealed by the lack of data on the populations of many species around the world (Hughes, 2017). Chiropterans are one of these taxa, as information on their populations is still poor despite the many threats they face (Furey & Racey, 2016; Mickleburgh et al., 2002).

Of the 1,500 recognised bat species worldwide, 38% are cave-dwelling species (IUCN, 2023; Simmons & Cirranello, 2025, Tanalgo et al., 2022), living in caves or natural or man-made underground cavities (e.g. mines or tunnels). These habitats are crucial for bats, which use them to reproduce, raise their young, rest on a daily or temporary basis and hibernate (Barros et al., 2020; Keeley & Tuttle, 1999; Kunz, 1982; Kunz et al., 2003; Lewis, 1995; Mering & Chambers, 2014; Ormsbee et al., 2007; Russo & Ancillotto, 2015; Struebig et al., 2009; The Protection of Bat Roost Guidelines Subcommittee et al., 1992). In tropical environments, most cave-dwelling species use caves as permanent roosts, and several species can co-occur in groups of hundreds or even thousands (Kunz, 1982; Monadjem et al., 2020). However, despite their ecological importance, these fragile environments are threatened and still poorly protected on a global scale (Balogh et al., 2020; Medellin et al., 2017; Neuhuber et al., 2020). Threats include resources extraction (minerals, guano, etc.) or recreational activities (caving, tourism, etc.) (Okonkwo et al., 2017; Simons, 1998). In turn, cave-dwelling bats generate ecological niches for diverse cave communities through the resources they produce (e.g., live and dead-bats, guano) (Labadie et al. 2025a).

Deleva et al. (2023) highlight the need to identify and protect key underground roosts. Broader global syntheses emphasize that effective conservation of bats roosting underground requires coordinated protection, monitoring, and policy measures (Meierhofer et al., 2024). Recent tools, such as the bat cave vulnerability index, provide integrative frameworks for assessing and ranking cave vulnerability at multiple scales (Tanalgo & Hughes, 2024). Unfortunately, due to largescale underfunding for bat research and conservation, the ecological information available on cave-dwelling bats is often too limited to inform these tools. In particular, data on bat populations and annual phenology of the occupation of caves is lacking, especially in African countries (Moutaouakil et al., 2024; Pretorius et al., 2021). Understanding when and how bats use caves throughout the year is therefore fundamental to assigning appropriate conservation value to each site. For example, a cave that hosts colonies during key reproductive periods (e.g. parturition or lactation) contributes critically to population persistence and should thus receive a higher conservation rank than a site used only for transient or non-reproductive activities.

The annual phenology of species at a given site is the result of the adaptation of its activity patterns to meet three primary needs: feeding, reproduction or rest (Alcock, 2013; Appel et al., 2019; Jones & Rydell, 1994). The phenology of bat activity in tropical caves reflects reproductive cycles adapted to local environmental conditions. In African regions, the timing of parturition in cave-dwelling insectivorous bats is closely linked to rainfall and subsequent insect availability (Cumming & Bernard, 1997). Insect abundance peaks about a month after peak rainfall, while births occur earlier to ensure weaning before maximum prey availability (Cumming & Bernard, 1997; Happold & Happold, 1990). This synchrony, though variable among species, highlights strong selective pressures imposed by seasonal food dynamics. Such cave-associated activity patterns are especially critical during maternity, when energetic demands are highest, and their alteration may signal adaptation to changing microclimatic or trophic conditions (Nkoana, 2020; Wolkovich et al., 2014).

Historically, bat communities have been studied using both invasive and non-invasive methods. Invasive methods, such as bat capture, remain often necessary to make an exhaustive species inventory, verify individual reproductive status and collect biometric data or samples (Appel et al., 2021; Barnhart & Gillam, 2014; Flaquer et al., 2007). Direct observation of individuals in their roost (e.g., cave, tree) or when they emerge to forage have also provided information on their behavior, abundance and movements (Thomas & West, 1989). Visual counts are sometimes difficult because of the large number of bats or the difficulty accessing the roosting sites. In addition, human observations are time-consuming and observer bias (Forsyth et al., 2006; Koger et al., 2023; Westcott & McKeown, 2004).

The development of new technologies have made it possible to set up precise counts of individuals in caves using digital image analysis, thermal cameras (Azmy et al., 2012; Sabol & Hudson, 1995) or bioacoustics (Sugai et al., 2020; Whiting et al., 2021). Passive acoustic monitoring (PAM) has a number of advantages: it is non-invasive, applicable in all environments (even the most remote ones), can collect a large amount of information such as abundance proxies, and the analysis can be automated (Hill et al., 2019). PAM exploits the use of echolocation to move and forage by more than 70% of the world’s bats (Browning et al., 2017; Gibb et al., 2019), Pteropodidae excluded. Recording these often species-specific sounds makes it possible to make a quantitative monitoring of bat species and/or communities. In recent years, several protocols, including that of Revilla-Martin et al (2020), have demonstrated the reliability of this technique compared to other methods for counting bat populations as they leave caves. Recent work by Appel et al. (2025) demonstrated that combining active and passive monitoring and optimally placing recorders at cave entrances maximizes species detection and comparability across sites and seasons.

However, PAM also faces limitations, such as the difficulty to distinguish between overlapping calls. This issue is compounded by the lack of reference call libraries in many regions, especially in Central Africa, and thus the impossibility to identify species in massive recordings. Sonotype classifiers were recently developped to counter this issue (Roemer et al., 2019) and opened the possibility to classify bat sounds in shapes and frequency groups where no species classifier is available. Labadie et al. (2025b) proposed a method to apply a sonotype classifier in association with local reference calls collected at cave entrances to obtain sound identification at the species level, or acoustic group, in massive recordings in the Republic of Congo. However, no application of this method has been reported so far.

The aim of this study was to demonstrate the possibility to conduct quantitative studies of cave-dwelling bat communities using acoustics in data-deficient areas. The Republic of Congo has very limited information on the acoustic repertoire of bats (Bates et al., 2013). Over a period of 19 months, we set up a PAM to describe the annual phenology of bat activity in two caves. We used the Tadarida sonotype classifier to classify bat vocalizations. To associate each sonotype with a species, we performed captures to obtain a species list and record reference calls, following the approach of Labadie et al. (2025b, c). To progress in the understanding of the biology and the ecology of bats in Central Africa, we compared seasonal variation in activity with the reproductive status of bats assessed from capture data. We expected species to show higher activity during periods of breeding and juvenile rearing (Catto et al., 1995; Dietz & Kalko, 2006).

## Methods

### Study area

Our study took place in the Republic of Congo, in a highly karstic region favouring the presence of numerous caves and/or cavities of various sizes in a rolling habitat of herbaceous savannah with patches of tropical forest. We selected two caves for the study: Boundou and Mont Belo, located in the Bouenza and Niari departments, about 50 km apart. Both caves are close to Dolisie, one of the three most important towns in the Republic of Congo on the southbound Brazzaville - Pointe Noire route. In this region, the seasons are divided into four: the short dry season (January to February), the short rainy season (March to May), the long dry season (June to mid-September) and the long rainy season (mid-September to December).

The Mont Belo Cave has several chambers (a main chamber and two secondary chambers). The Boundou Cave, on the other hand, is a smaller cavity integrated into a rocky complex, with a main chamber and an annex chamber that is very difficult for humans to access, and houses a small internal lake. Details of the configuration of the two caves in this study are presented in Fig. 1.

**Figure 1.**
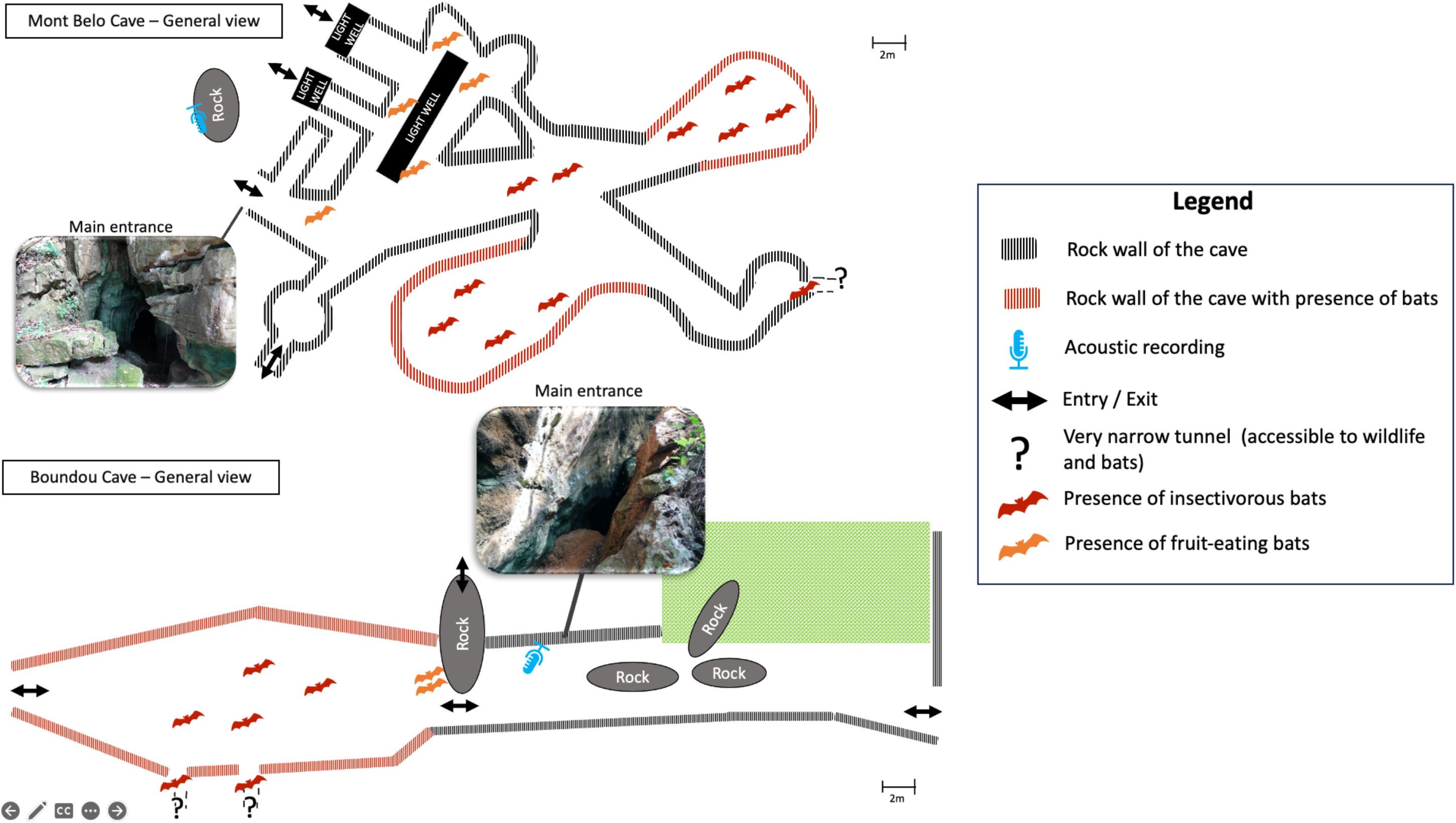
General diagram of the configuration of the Boundou and Mont Belo Cave, the location of the acoustic recorder and the bat roosting sites.

### Ethic statements

The protocols presented below have all been validated by the Ethics Committee of the Ministry of Scientific Research and Technological Innovation and by the Ministry of Forest Economy of the Republic of Congo (N°212/MRSIT/IRSSA/CERSSA and N°687/MEF/CAB/DGEF-DFAP). This work was also carried out with the agreement and participation of the authorities in the Niari and Bouenza departments, as well as the communities and landowners close to our study sites.

### Capturing bats

During the 19 months of data collection (September 2021 – March 2023), we conducted eleven capture sessions at the main entrance of each cave using a harp trap (© Ecotone) between 5:00 and 10:00 pm. The sessions were distributed across different months and seasons to capture temporal variation in activity and reproductive condition. However, sampling did not systematically cover all months of the annual cycle at each site. A detailed summary of sampling dates, months covered, and numbers of individuals captured per species and reproductive category is provided in Table 1 and Appendix 1. During the first session at Mont Belo in September 2021, no bats were captured, likely due to a nearby crop fire that may have forced bats to temporarily abandon the cave. In total, 672 bats were captured, including 344 individuals at Boundou cave and 328 at Mont Belo cave. During these capture sessions, a wide range of information was collected, including the time of capture, the family and species identified in the field, sex, age and reproductive status. The reproductive status of individuals was determined by assessing females for pregnancy (abdominal palpation) and lactation (mammary gland size, absence of fur, presence of milk at the nipple). Males were assessed for sexual activity based on testes size and epididymal coloration. At the beginning of the study, in order to confirm the genera and species of bats present in the caves, we collected genetic samples, acoustic recordings and specimens for reference purposes.

**Table 1:** Total number of individuals captured by species, cave, and sex, with observed reproductive periods, confidence levels, and capture sessions (September 2021–March 2023). Female reproductive periods include gestation, parturition, and lactation. Male sexual activity inferred from testes size and epididymal coloration. Confidence level (Conf. level) symbols are automatically derived from the raw data by acoustic group × cave. ‘?’ = insufficient data. See footnote for legend and species notes.

| Group | Species | Cave | Males | Females | Total | Female reproductive period(s) | Conf. level | Male sexual activity | Conf. level | Capture sessions (month, year) |
| --- | --- | --- | --- | --- | --- | --- | --- | --- | --- | --- |
| *Hipposideros caffer* | *Hipposideros cf. caffer* | Mont Belo | 62 | 17 | 79 | Jan, Apr, Oct, Dec | (S) | Jan, Feb, Oct | (I) | Oct 2021, Dec 2021, Feb 2022, Apr 2022, Jun 2022, Sep 2022, Oct 2022, Nov 2022, Jan 2023, Mar 2023 |
|  | *Hipposideros cf. caffer* | Boundou | 43 | 82 | 125 | Feb, Jun, Sep, Oct, Nov, Dec | (C) | Jan, Mar, Jun, Sep, Oct, Nov | (S) | Sep 2021, Oct 2021, Dec 2021, Feb 2022, Apr 2022, Jun 2022, Sep 2022, Oct 2022, Nov 2022, Jan 2023, Mar 2023 |
| *Miniopterus group* | *Miniopterus cf. minor* | Boundou | 41 | 25 | 66 | Mar, Sep, Oct | (C) | Jan, Jun | (I) | Sep 2021, Oct 2021, Dec 2021, Feb 2022, Apr 2022, Jun 2022, Sep 2022, Oct 2022, Jan 2023, Mar 2023 |
|  | *Miniopterus cf. minor* | Mont Belo | 14 | 18 | 32 | Jun, Sep, Oct, Nov | (C) | Jun, Dec | (I) | Oct 2021, Dec 2021, Feb 2022, Jun 2022, Sep 2022, Oct 2022, Nov 2022, Jan 2023, Mar 2023 |
|  | *Miniopterus cf. inflatus* | Boundou | 2 | 1 | 3 | Jun | (C) | ? | (I) | Apr 2022, Jun 2022 |
|  | *Miniopterus cf. inflatus* | Mont Belo | 77 | 25 | 102 | Apr, Jun, Sep, Oct | (C) | Nov | (I) | Oct 2021, Dec 2021, Feb 2022, Apr 2022, Jun 2022, Sep 2022, Oct 2022, Nov 2022, Jan 2023, Mar 2023 |
| *Rhinolophus denti* | *Rhinolophus cf. denti* | Boundou | 4 | 2 | 6 | Oct | (S) | ? | (I) | Dec 2021, Feb 2022, Oct 2022, Nov 2022 |
|  | *Rhinolophus cf. denti* | Mont Belo | 31 | 71 | 102 | Jan, Apr, Jun, Sep, Oct, Nov, Dec | (C) | Feb, Nov | (I) | Oct 2021, Dec 2021, Feb 2022, Apr 2022, Jun 2022, Sep 2022, Oct 2022, Nov 2022, Jan 2023, Mar 2023 |
| *Rhinolophus landeri-alcyone* | *Rhinolophus cf. landeri-alcyone* | Mont Belo | 9 | 4 | 13 | Jan, Oct, Nov | (S) | ? | (I) | Oct 2021, Feb 2022, Apr 2022, Oct 2022, Nov 2022, Jan 2023, Mar 2023 |
| *Rhinolophus Macro Tria Group* | *Rhinolophus cf. landeri-alcyone* | Boundou | 6 | 33 | 39 | Jan, Oct, Nov | (C) | ? | (I) | Oct 2021, Dec 2021, Apr 2022, Oct 2022, Nov 2022, Jan 2023, Mar 2023 |
|  | *Triaenops cf. afer\** | Boundou | 42 | 58 | 100 | Jan, Feb, Apr, Jun, Sep, Oct, Nov, Dec | (C) | Nov | (I) | Sep 2021, Oct 2021, Dec 2021, Feb 2022, Apr 2022, Jun 2022, Sep 2022, Oct 2022, Nov 2022, Jan 2023, Mar 2023 |
|  | *Macronycteris cf. gigas\** | Boundou | 3 | 1 | 4 | Nov | (C) | ? | (I) | Dec 2021, Apr 2022, Nov 2022 |

\* Species found only in the Boundou cave. Confidence level: (C) Confirmed: ≥ 5 females AND ≥ 2 positive reproductive indicators in at least one month; (S) Suggested: ≥1 positive indicator (pregnant, lactating, or carrying young); (I) Insufficient: 0 positive indicators.

### Passive acoustic monitoring (PAM)

At each main entrance of the two caves, at a maximum distance of five meters, we placed an SM4BAT acoustic recorder with a U1 microphone (Wildlife acoustics, Maynard, MA, USA). Each microphone was placed on a pole or tree and oriented to best record the acoustic calls of bats emerging and re-entering through the main entrance. These recorders ran continuously several weeks per month for a period of 19 months (September 2021 to first day of March 2023). The SM4BATs were set up according to the stationary points protocol of Vigie-chiro, a citizen science programme for the acoustic monitoring of bat populations in France (French Museum of Natural History, see Mariton et al., 2023) starting 30 minutes before sunset and 30 minutes after sunrise. Figure 1 show the location of the acoustic recorder in relation to the configuration of each cave.

### Processing acoustic data

Recordings were cut in durations of five seconds maximum. In order to process all the acoustic files collected during this study in the two caves, we used the automatic sonotype classifier published in Roemer et al., (2021) and modified in Labadie et al. (2025b) to identify the presence of different sonotypes inside each file. The classifier returns a table in which each line corresponds to a group of calls with similar call shape and frequency, associated with information about their acoustic parameters (e.g., frequency, duration). Because of an overlap in the shape and frequency of certain acoustic calls from bat species of the same family, we decided to group the species at each study site using a posteriori classification (detail for the different acoustic groups in Labadie et al., 2025b). The Boundou cave has four distinct groups: (1) the *Miniopterus* group (*Miniopterus cf minor*/ *Miniopterus cf. inflatus*), (2) *Hipposideros caffer* (*Hipposideros cf. caffer)*, (3) *Rhinolophus denti* (*Rhinolophus cf. denti*) and (4) the *Rhinolophus* - Macro - Tria group (*R. cf. landeri-alcyone* /*Macronycteris gigas*/*Triaenops cf. afer*) (Fig. 2). The Mont Belo cave also has four distinct groups: (1) the *Miniopterus* group (*M. cf minor*/ *M. cf. inflatus*), (2) *Hipposideros caffer* (*Hipposideros cf. caffer*), (3) *Rhinolophus denti* (*Rhinolophus cf. denti)* and (4) *Rhinolophus landeri-alcyone (Rhinolophus cf. landeri-alcyone*) (Fig. 2). For more details on the automatic classification and on a posteriori sorting of acoustic groups, please respectively refer to Roemer et al. (2021) and Labadie et al. (2025b).

**Figure 2.**
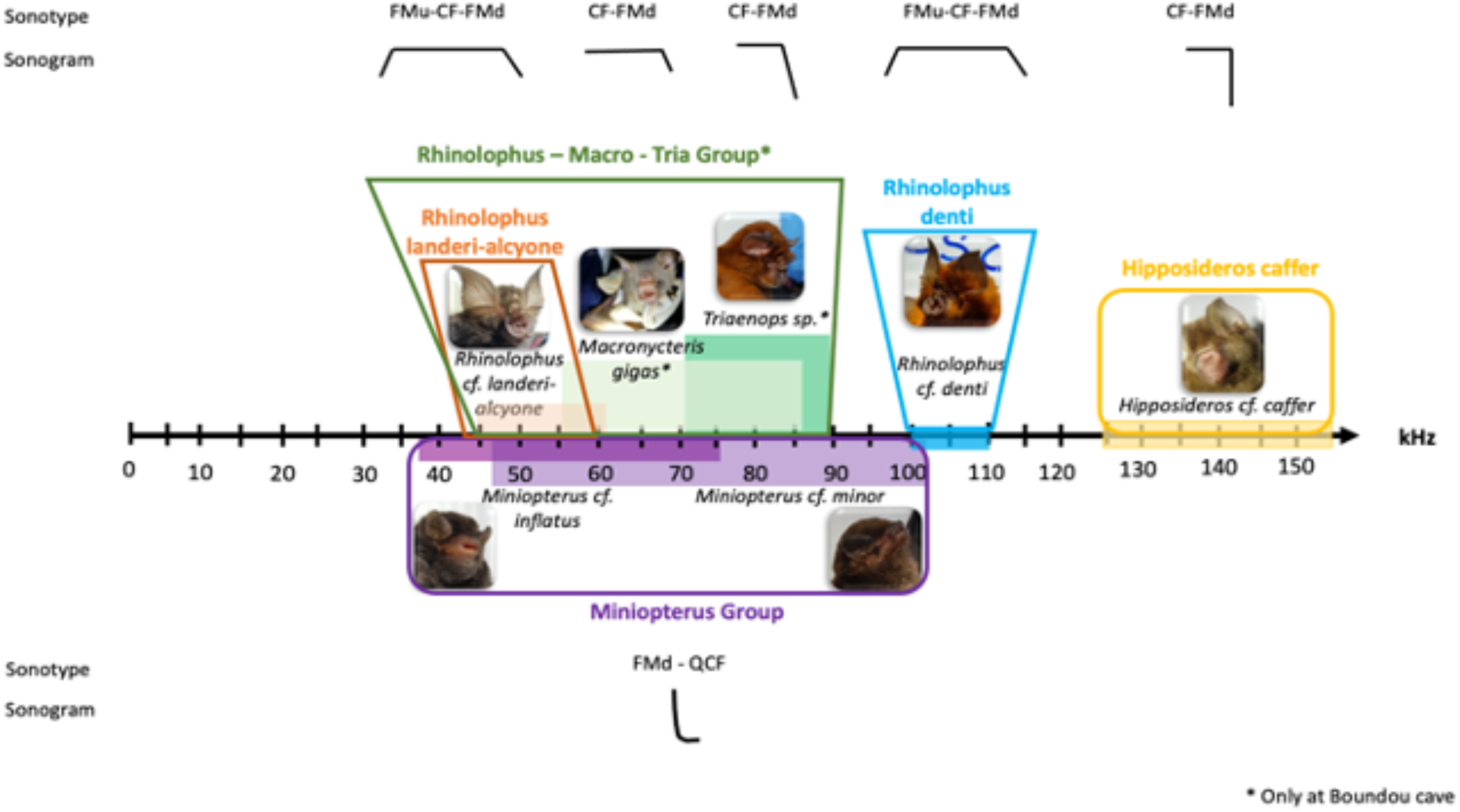
Acoustic characteristics for the different bat species captured at the two study sites and categorization into different acoustic groups or species. Detail of sonotype, sonogram and acoustic frequency recorded or listed in reference works for each bat genus and species captured in the two caves (species with * are present only at Boundou). In the Boundou cave, there are four acoustic groups (*Rhinolophus* – Macro – Tria group; *Rhinolophus denti*; *Hipposideros caffer* and *Miniopterus* group) and in the Mont Belo cave there are four acoustic groups (*Rhinolophus landeri-alcyone*; *Rhinolophus denti*; *Hipposideros caffer* and *Miniopterus* group).

### Statistical analysis

After analyzing all recorded files, the Tadarida acoustic classifier produced raw results, which were then organized in the post-sorting output. All the statistical analyses presented below were carried out using R software v. 4.3.1 (R Core Team, 2023). The graphics were produced using ggplot2 (Wickham, 2016) or ggradar (Bion, 2023).

### Bat counting techniques and acoustics analyses

Bats may emerge from and enter a cave at any time during the night; nonetheless, most of them emerge after sunset and enter before sunrise (Erkert, 1982). In order to assess the activity of the five groups of bats during emergence and their entry at each cave during the 19 months, we decided to standardise our methodology based on the protocol of Mariton et al. (2023). As in other acoustic studies (Millon et al., 2015), we defined the activity of the different groups of bats as the number of bat passes. A bat pass is the occurrence of a single or several calls detected during a five seconds interval.

As our study took place in the tropics and in two caves that are geographically close, the characterisation of the diel activity of groups of bats can be based on a time scale (on sunrise and sunset). To begin with, we defined the time of sunset and sunrise based on the geographical coordinates and using the getSunlightTimes function in the suncalc package (Thieurmel et al., 2022). We then quantified the number of bat passes, over two specific time periods: (a) the period of emergence, from 30 minutes before sunset to two hours after sunset, and (b) the period of entry into the cave, from two hours before sunrise to 30 minutes after sunrise. This calculation was carried out for each species, each site and each day on which a complete night was recorded by the SM4BATs.

### Distribution of activity during the night

To calculate the distribution of the activity of acoustic groups of bats throughout the night, we estimated the activity density using the density function in the R package. This calculation was done for the twelve months of the year (some months were sampled repetitively during different years) for each species and each study site.

### Monthly pattern of the emergence and the entry activity at the roost

We used Generalised Linear Mixed Models (GLMM) to model the number of bat passes (proxy of activity) during their emergence and entry as a function of the month of the year in each cave and for each acoustic group. We added month and year as a random effect (1|month_year). In total, we fitted sevent separate emergence models (see Appendix 2). Each model corresponds to a specific bat species or acoustic group at a given cave. This approach allows us to account for species-specific emergence patterns at each site while controlling for temporal variation across months and years. We used the glmmTMB package (Brooks et al., 2017) to build our models. We used a lognormal distribution and checked the conditions and diagnostics using the DHARMa package (QQ plot residuals, Residuals vs predicted, dispersion, zero-inflation) (Hartig & Hartig, 2022). We used a Bonferroni correction to limit the type I error rate (false positive) when interpreting generalised linear models (Cabin & Mitchell, 2000).

## Results

### Species composition and reproductive phenology

In both caves, we captured a total of seven bat species. Five species were present at both sites: *Miniopterus cf. minor* (N = 98), *Miniopterus cf. inflatus* (N = 105), *Rhinolophus cf. landeri-alcyone* (N = 52), *Rhinolophus cf. denti* (N = 108), and *Hipposideros cf. caffer* (N = 204). *Triaenops cf. afer* (N = 100) and *Macronycteris cf. gigas* (N = 4) were recorded exclusively in the Boundou Cave. For each species, the number of captured individuals by sex and by site is summarized in Table 1, together with information on reproductive status.

Across species and sites, most bats exhibited a female reproductive period (gestation, parturition, and lactation) during the long rainy season (October–December). Male sexual activity was mainly observed during the long dry season (June–mid-September), suggesting a seasonal separation between mating and juvenile rearing (Table 1). Some taxa, particularly *Hipposideros cf. caffer*, showed evidence of two reproductive periods within the annual cycle.

### Passive acoustic monitoring

In total, during nineteen months, we collected 2,375,956 acoustic files (five seconds duration) with 48.7% coming from Boundou and 51.3% from Mont Belo. The acoustic groups identified at each cave are shown in Figure 2 (Appendix 1, raw data for each group).

### Nocturnal activity patterns

In both caves, for the five acoustic groups (*Hipposideros caffer, Miniopterus group, Rhinolophus denti, Rhinolophus landeri-alcyone, Rhinolophus Macro Tria group*), nocturnal activity patterns, specifically emergence and return activity, were generally bi-modal. More precisely, the first peak of activity occurred at the beginning of the night, around 6:00 pm (after sunset), and a second peak was observed between 5:00 and 6:00 am, depending on the species (Fig. 3 and Fig. 4). In contrast, Rhinolophus cf. denti showed an earlier onset of activity, beginning around 3:00 am in both caves.

**Figure 3.**
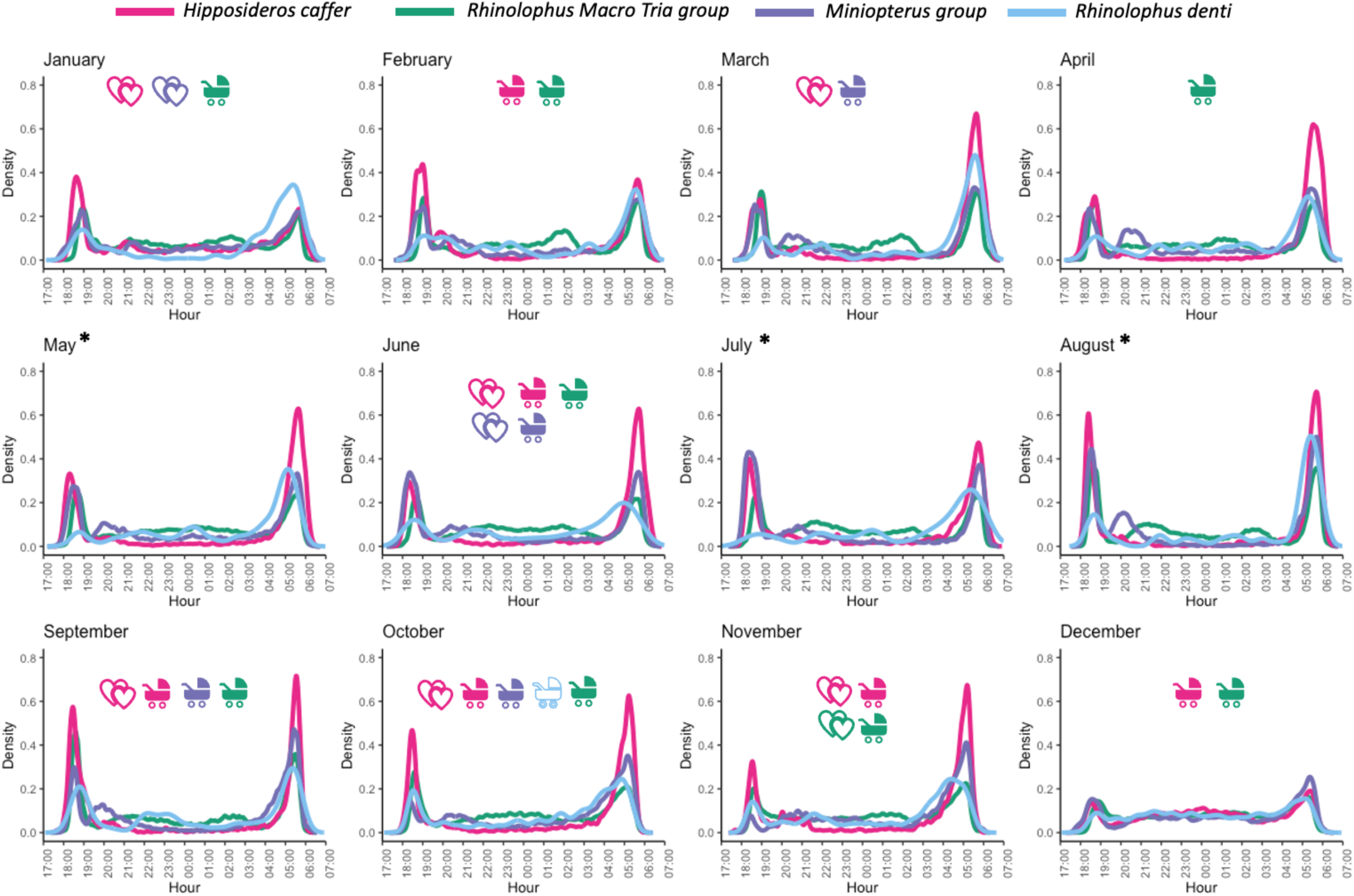
Number of bat passes for each acoustic group according to month at Boundou cave. The reproductive status of the species or acoustic group is indicated by two pictograms with the color of the acoustics groups: 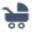 female reproductive period (lactation, pregnant, young rearing) and 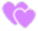: mating period/male activity. The months of May, July and August, which have an *, are month when no captures were made, hence the absence of data on the reproductive status of the acoustic group. Outlined icons indicate a suggested (S) or insufficient (I) confidence level; filled icons indicate confirmed (C).

**Figure 4.**
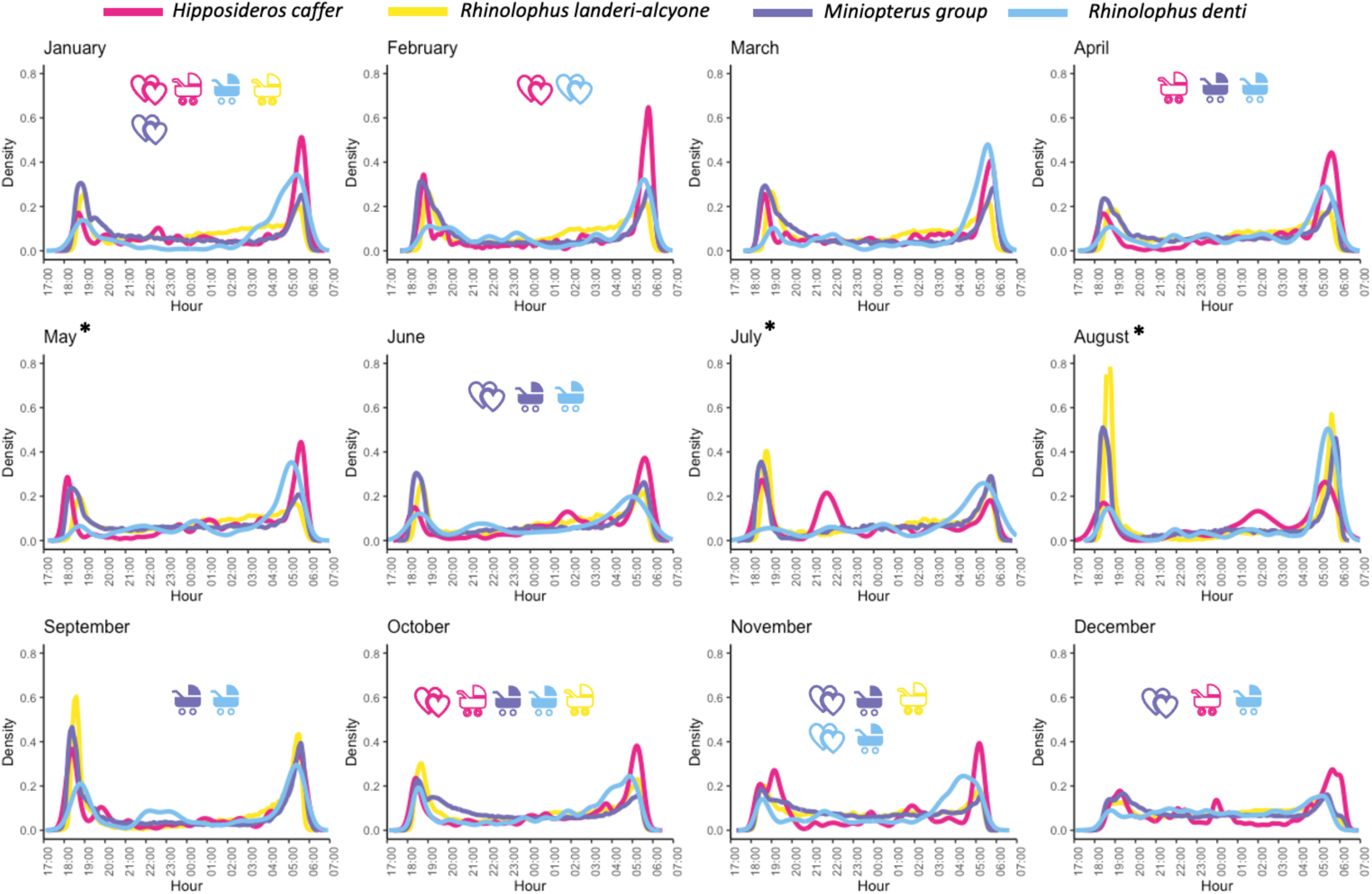
Number of bat passes for each acoustic group according to month at Mont Belo cave. The reproductive status of the species or acoustic group is indicated by two pictograms with the color of the acoustics groups: 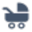 female reproductive period (lactation, pregnant, young rearing) and 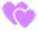: mating period/male activity. The months of May, July and August, which have an *, are month when no captures were made, hence the absence of data on the reproductive status of the acoustic group. Outlined icons indicate a suggested (S) or insufficient (I) confidence level; filled icons indicate confirmed (C).

In December, bat activity was more evenly distributed throughout the night than during the other months for all five acoustic groups and in both caves (Fig. 3 and Fig. 4). In addition, the Boundou cave activity was more spread-out during January and February (Fig. 3). In the Mont Belo cave, activity was also more spread out throughout the night in November and July (Fig. 4).

### Annual phenology of bat activity and predicted species-specific patterns

Because analyses using emergence and entry activity produced similar annual phenology of bat activity, we decided to comment our results for the emergence activity only (Fig. 5) (Appendix 4, 5, 6 and 7 for entry activity prediction). The annual phenology of activity varied according to the acoustic group and the month (Appendix 2 and 3). For *Rhinolophus denti*, their low activity during emergence in Mont Belo cave meant that the model had no results (the model had too little data to run) (Appendix 6 and 7).

**Figure 5.**
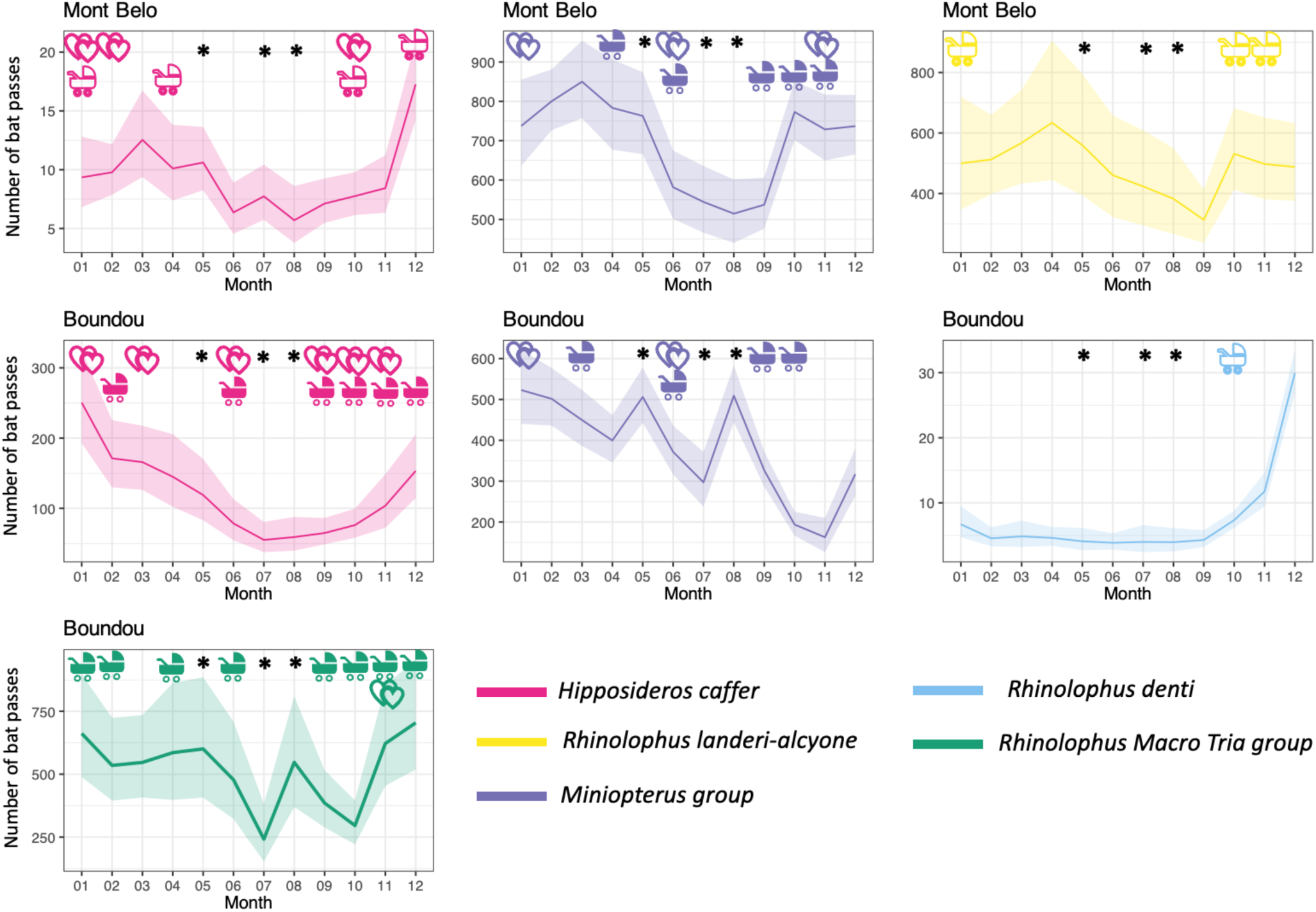
Prediction of acoustic group emergence activity (estimated based on the number of bat passes) with 95% confidence interval as a function of months and study sites (Boundou and Mont Belo cave). The *Rhinolophus dent*i group at Mont Belo has not been represented here due to lack of data. The reproductive status of the species or acoustic group is indicated by two pictograms with the color of the acoustics groups: 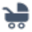 female reproductive period (lactation, pregnant, young rearing) and 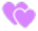 : mating period/male activity. The months of May, July and August, which have an *, are month when no captures were made, hence the absence of data on the reproductive status of the acoustic group. Outlined icons indicate a suggested (S) or insufficient (I) confidence level; filled icons indicate confirmed (C).

#### Hipposideros caffer

Predicted bat activity varied seasonally across species groups and caves (Figs. 4 and 5). For *Hipposideros caffer*, activity was generally moderate during the short dry (January–February) and short rainy seasons (March–May) in both caves. Nightly activity patterns (Fig. 4) were consistently bimodal, with distinct emergence and entry peaks. During the long dry season (June–August), activity declined, particularly at Boundou. Activity increased again toward the long rainy season (September–December), when nightly activity became more evenly distributed across the night, coinciding with the main juvenile-rearing period. According to Fig. 5 (Appendix 8 predicted activity remained relatively stable at Mont Belo from January to April, while Boundou showed higher values during the same period. The highest predicted activity occurred at the end of the year in both caves.

#### Miniopterus group

The Miniopterus group showed the highest activity during the short dry and short rainy seasons in both caves (Fig. 4). During the long dry season, activity declined and nightly patterns became less structured. Activity remained lower during the long rainy season, despite the presence of juveniles, with reduced intensity of emergence and entry peaks. Fig. 5 (Appendix 8) indicates that predicted activity peaked between January and March at Mont Belo and remained high from January to May at Boundou. Minimum activity was observed during the long dry season, especially in July–August.

#### Rhinolophus landeri-alcyone

At Mont Belo, *Rhinolophus landeri–alcyone* exhibited low activity during the short dry season, followed by an increase during the short rainy season (Fig. 4). Activity decreased markedly during the long dry season, when nightly activity was minimal. An increase was observed again during the long rainy season, corresponding to pregnancy, parturition, and lactation, with more continuous nocturnal activity. As shown in Fig. 5 (Appendix 8), predicted activity reached its lowest values between July and September and increased again from October to December. The highest monthly values occurred during the late rainy season.

#### Rhinolophus denti

At Boundou, *Rhinolophus denti* showed low activity for most of the year, particularly from the short rainy season through the long dry season. A clear increase occurred during the long rainy season, with the highest activity at the end of the year (Fig. 4). Fig. 5 (Appendix 8) shows consistently low predicted activity from February to September, followed by a marked increase from October to December. The end of the year corresponds to the period of highest predicted bat passes.

#### Rhinolophus Macro-Tria group

Finally, the *Rhinolophus Macro–Tria group* at Boundou showed higher activity during the short dry and short rainy seasons (Fig. 4). Activity declined during the long dry season and remained generally low during the long rainy season, with reduced nightly structure. According to Fig. 5 (Appendix 8), predicted activity was highest in January–February and declined sharply during the long dry season. A secondary increase was observed at the end of the year.

## Discussion

We used passive acoustic monitoring to characterize bat communities and their phenologies at the entrances of caves in the Republic of Congo, addressing key challenges in cave-dwelling bat research in data_deficient countries. A global sonotype classifier augmented with locally recorded reference calls allowed to overcome key limitations in these contexts : the absence of pre-existing reference acoustic libraries, the difficulty of monitoring nocturnal and cryptic species in remote places and the absence of standardized frameworks for quantifying bat activity. In two caves, we quantified the activity of five acoustic groups of bats and described their nightly and annual phenologies. This methodological adaptation combining sonotypes with on-site reference recordings offers a framework that can be replicated in other regions lacking acoustic data, potentially across different biomes and bat communities.

### Nocturnal activity pattern

Five acoustic groups were detected in both caves: *Hipposideros caffer*, the *Miniopterus* group, *Rhinolophus denti*, *Rhinolophus landeri-alcyone*, and the *Rhinolophus Macro Tria group*. Overall, their emergence and entry patterns showed similar dynamics, with two bi-modal nocturnal activity patterns (a period of emergence and a period of entry into the cave) for most months in both caves. In several months, however, nocturnal activity patterns was more evenly distributed throughout the night—particularly in December for all groups, in January–February at Boundou, and in November and July at Mont Belo. When outside the cave, the large number of individuals observed are hypothesized to go hunting before returning to the cave. Some individuals may move to another resting place or return later.

Nightly observations revealed patterns consistent with known ecological behaviors. With a few exceptions, bats emerged from their daytime roosts at dusk—around 6 p.m.—to fly to feeding sites, as expected (Bullock et al., 1987). Emergence timing was correlated with the time of sunset (Erkert, 1982). A second peak of nocturnal activity generally occurred during the early morning hours, around 5–6 a.m., corresponding to their return to the caves. Central African countries such as Congo have little variation in sunrise and sunset throughout the year (average sunrise: about 05:50 – 06:02 am local time; average sunset: about 05:55 – 06:05 pm local time) (WorldData, 2024), leading to limited variation in emergence timing. In our study, we also observed “burst activity” patterns during emergence, where several bats exited the cave in rapid succession followed by short periods of reduced activity, a behavior previously described in other bat colonies (Swift, 1980). This more even distribution of nightly activity during October–January coincided with periods when pregnant and lactating females were recorded, suggesting that reproductive constraints may influence activity patterns. However, given the uncertainty associated with the exact timing and duration of reproductive periods, this relationship should be interpreted with caution. These nocturnal activity patterns provide the detailed temporal resolution necessary to interpret the annual phenology of bat activity, which is strongly shaped by reproductive cycles and resource availability.

### Annual phenology of bat activity and reproductive period

Building on nocturnal activity patterns, we examined the annual phenology of bat activity across the year to understand seasonal trends. In our study, activity peaks during the main rainy season (October–December) coinciding with maternity colony formation (gestation, birth, and beginning of lactation). While this temporal overlap is consistent with reproductive activity, the precise boundaries of these periods remain uncertain and may vary among species and sites. In December and January, emergence nocturnal activity was spread throughout the night likely reflecting females alternating between foraging and suckling their young, as reported in previous studies (Catto et al., 1995; Dietz & Kalko, 2006; Maier, 1992; Swift, 1980, 1997). Field observations support this interpretation, suggesting repeated foraging–nursing movements between the cave and feeding sites. This pattern is consistent with Henry et al. (2002), who showed that *Myotis lucifugus* reduced their foraging range and returned to the roost one to two times per night. Such returns likely generate secondary activity peaks after initial emergence.

Our results also show peaks during the short rainy season (March–May), likely reflecting the independence of juveniles. The timing of independence varies among species. For example, young *Hipposideros caffer* become independent between three and three and a half months after birth (Wright, 2009). However, the young are not carried by their mother throughout this period, and may begin to fly one month after birth, staying in the roost with other individuals of the same generation when their mothers forage out of the cave (Wright, 2009).

Finally, based on capture observations, we hypothesise that the annual phenology of bat activity peaks observed during the dry season (January–February) correspond to the initiation of reproductive cycles for certain species, such as *Hipposideros caffer* and the *Miniopterus group*.

Some acoustic groups in our study showed two reproductive periods within the year. For example, the *Miniopterus* group exhibited peaks of reproductive activity in March–April and again from September to December, while *Rhino Macro Tria group* showed similar patterns between September–January and March–June. These patterns likely reflect the presence of multiple species within each acoustic group. These likely reflect species-specific reproductive strategies. In the case of *Hipposideros caffer*, the dual activity could be explained by the presence of a cryptic species complex within this acoustic group, as suggested in previous studies (Wright, 2009). Alternatively, the observed extended annual phenology of bat activity could reflect a somewhat flexible reproductive period. Although individual females may start reproduction at slightly different times, parturition generally occurs during the peak of insect availability in the rainy season. This partial synchrony allows offspring to benefit from abundant food resources while maintaining flexibility in reproductive timing, which can maximize survival (Racey & Entwistle, 2000; McWilliam, 1988). Such flexibility, combined with uncertainty in the exact timing of reproductive events, may contribute broad and overlapping activity patterns. This likely aligns activity with seasonal resource peaks.

The long dry season (June–August) showed reduced nocturnal activity patterns compared with the other seasons. This suggests that bats congregate less in these habitats due to limited resources, a pattern that aligns with previous studies (Barclay & Kurta, 2007; Patriquin & Ratcliffe, 2016; Patterson et al., 2007; Postawa & Gas, 2009). Scattered nocturnal activity patterns were observed throughout the night during the long dry season (June - August). Unfortunately, we did not capture in the months of July and August, and could not assess the reproductive status of the individuals. However, the gestation period observed from late September to early December in all acoustic groups suggests that copulation likely occurs towards the end of the long dry season (June–July). This timing aligns with swarming behavior and is consistent with observations from other countries. For example, *Hipposideros caffer* typically mates at the end of the first rainy season (June–July) and gives birth during the middle of the second rainy season (October–November) (Bernard, 1982; Brosset, 1968; Günther et al., 2016).

The only exception was the *Miniopterus* group in Boundou cave, which remained very active in August (long dry season). The females of this species may use the cave to gather in a maternity colony before giving birth in September and October (ML personal observation). This is consistent with Cumming and Bernard (1997) who mentioned that many bat species give birth to their young about a month before the peak in resource availability.

Finally, variation in roost use by species and cave highlights the complexity of bat ecology in these systems. *Rhinolophus denti* appeared to use caves solely for rearing young, not for mating. Conversely, *Hipposideros caffer* used both caves for mating and rearing young. It is noteworthy that the mating period of *Hipposideros caffer* seemed to vary between cave (January, February, June in Mont Belo cave and March and June in Boundou cave). This variation may be explained by the presence of two cryptic species of *Hipposideros* in this study (ML personal observation).

### Conservation implications for the caves and application in other biomes

By describing nocturnal activity patterns and annual phenology of bat activity, our results also highlight the ecological and conservation significance of these caves. Although this study did not aim to formally calculate the Bat Cave Vulnerability Index (BCVI) (from Tanalgo & Hughes, 2024), a quick assessment of standard criteria suggests differing vulnerability between the two caves. Mont Belo hosts more species and several maternity colonies. It therefore has the highest biotic value. Its vulnerability score must then be assessed by considering the different threats affecting the site (Labadie et al., 2025a). Boundou shows lower vulnerability, with fewer disturbances and only occasional predation (Labadie et al., 2025a).

### Limitation of this study

Information on reproductive status cannot be obtained from acoustic monitoring, it requires the capture of individuals. We were not able to capture during the long dry season (June–August), which hampers our understanding of reproductive status of the different species. More broadly, uncertainty in the timing and duration of reproductive periods limits the precision of phenological interpretations. We could not validate acoustic data with complementary methods (e.g., counts or video monitoring) due to logistical constraints. These complementary methods would have provided direct estimates of colony size and temporal fluctuations in bat abundance. Such information would have allowed us to link nocturnal activity patterns more precisely to actual population dynamics and to better distinguish between changes in the number of individuals. Acoustic activity is, however, generally correlated with colony size in caves (Revilla-Martín et al., 2020). Still, variations in flight behavior, such as increased vocal activity during social interactions or reduced calling during certain foraging strategies, may lead to an over- or underestimation of this correlation. Furthermore, for several species-site combinations, the confidence level assigned to reproductive periods was rated as suggested (S) or insufficient (I), reflecting limited sample sizes or a low number of positive reproductive indicators in a given month. These observations should be interpreted as preliminary signals rather than confirmed reproductive events, and targeted sampling efforts during these periods are needed to strengthen conclusions.

## Conclusion

We characterized the variation in activity of acoustic groups in relation to season and breeding status, providing insights into temporal use of roosts and potential sensitivity to disturbance. While both caves host similar species groups and share general seasonal trends, Mont Belo shows more distinct biannual reproductive peaks (especially for Miniopterus and R. landeri–alcyone), whereas Boundou displays more evenly distributed activity and possibly more stable year-round occupation (notably for *Hipposideros caffer* and the *Rhinolohus Macro Tria group*). These patterns suggest functional complementarity between the two caves, Mont Belo cave likely serving as a breeding and maternity site, and Boundou cave as a multi-species roost used across seasons. They also highlight the potential of PAM combined with acoustic classification to study reproductive behavior in wild populations.

Our study demonstrates that quantitative bat monitoring is possible even in data-deficient areas. This approach offers a valuable tool for cave conservation, as it enables the identification of critical periods of bat activity, reproductive timing, and species-specific cave use. Even with some uncertainty in reproductive phenology, this information is essential for prioritizing protection and managing disturbance of cave bats (Tanalgo & Hughes, 2024). A major strength of this study lies in its non-invasive approach, applicable across multiple caves and species, while obtaining both nocturnal activity patterns and annual phenology of bat activity.

Future improvements could include non-invasive hormonal monitoring by collecting bat urine to analyze stress levels and reproductive hormones, allowing identification of mating and lactation periods without capturing bats (Greville et al., 2022; Kumar & Umapathy, 2019; Voigt & Schwarzenberger, 2008; Wilkinson et al., 2024). Given the importance of bats for the health of the environment across the world, this study provides new avenues to produce knowledge in data-deficient and remote areas in order to adapt bat conservation and management.

## Appendices

**Appendix 1 Morphometric data of insectivorous bat communities in two caves of the Republic of Congo and acoustic data**

CIRAD Dataverse : morphometric data : https://doi.org/doi:10.18167/DVN1/M2J0TY.

Zotero : Acoustic data : https://doi.org/10.5281/zenodo.15636144

**Appendix 2.**
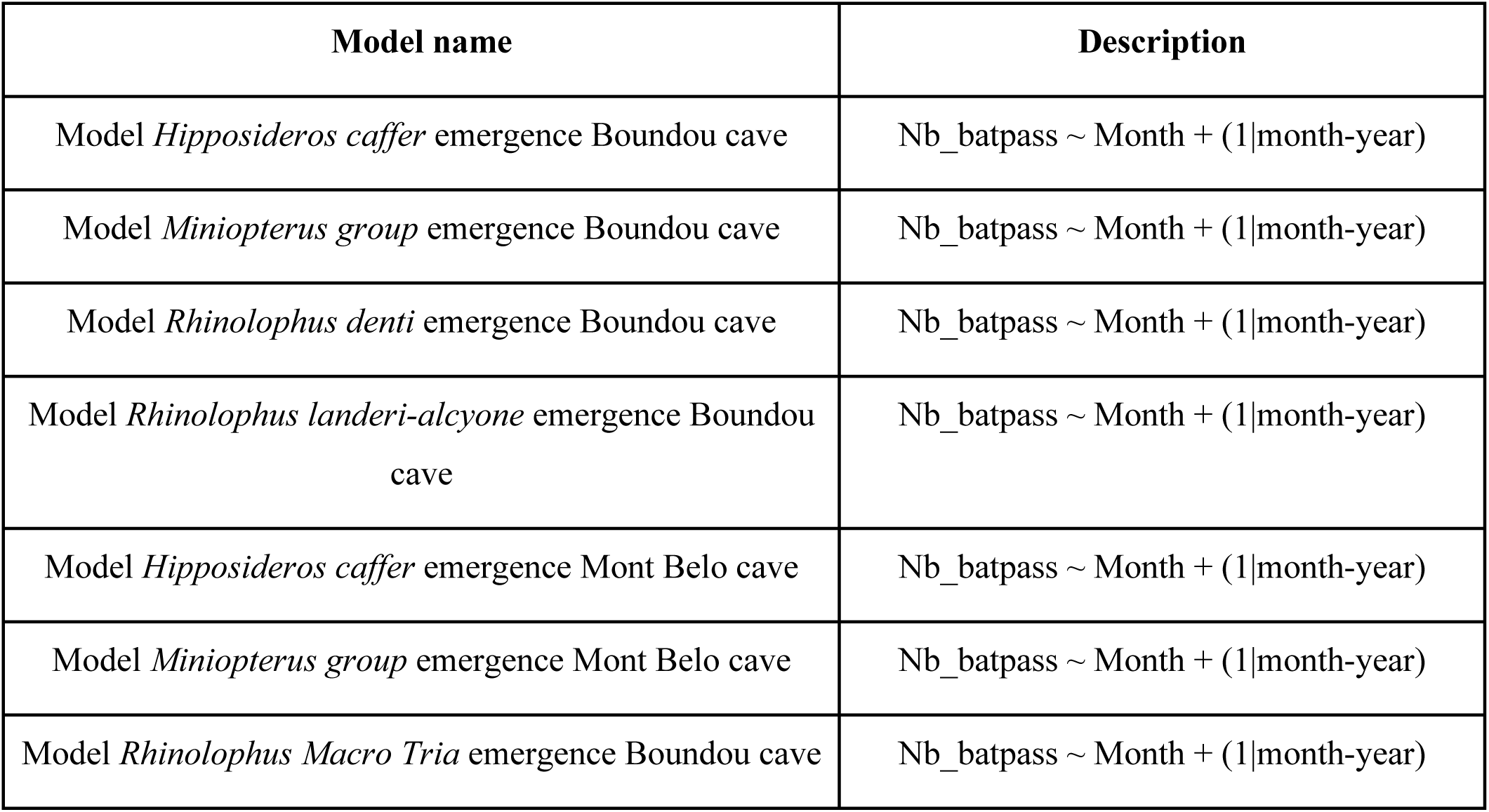
Summary table of seven emergence models for each cave (Boundou and Mont Belo) and each acoustics group tested in this study.

**Appendix 3.**
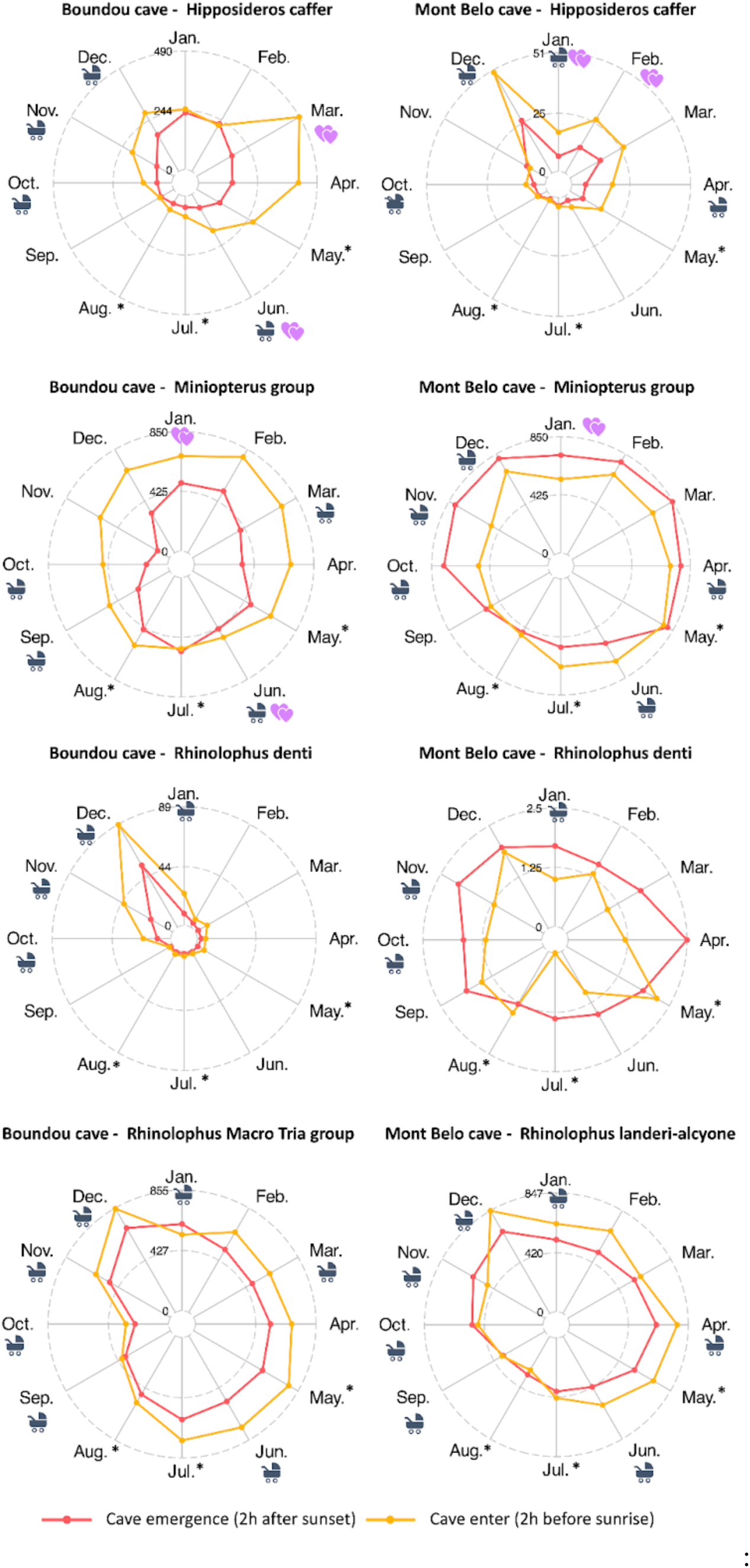
Number of bat passes for each acoustic group according to month and site (Boundou and Mont Belo Cave). The reproductive status of the species or acoustic group is indicated by two pictograms: 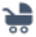 female reproductive period (lactation, pregnant, young rearing) and 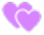: mating period/male activity. The months of May, July and August, which have an *, are month when no captures were made, hence the absence of data on the reproductive status of the acoustic group.

**Appendix 4.**
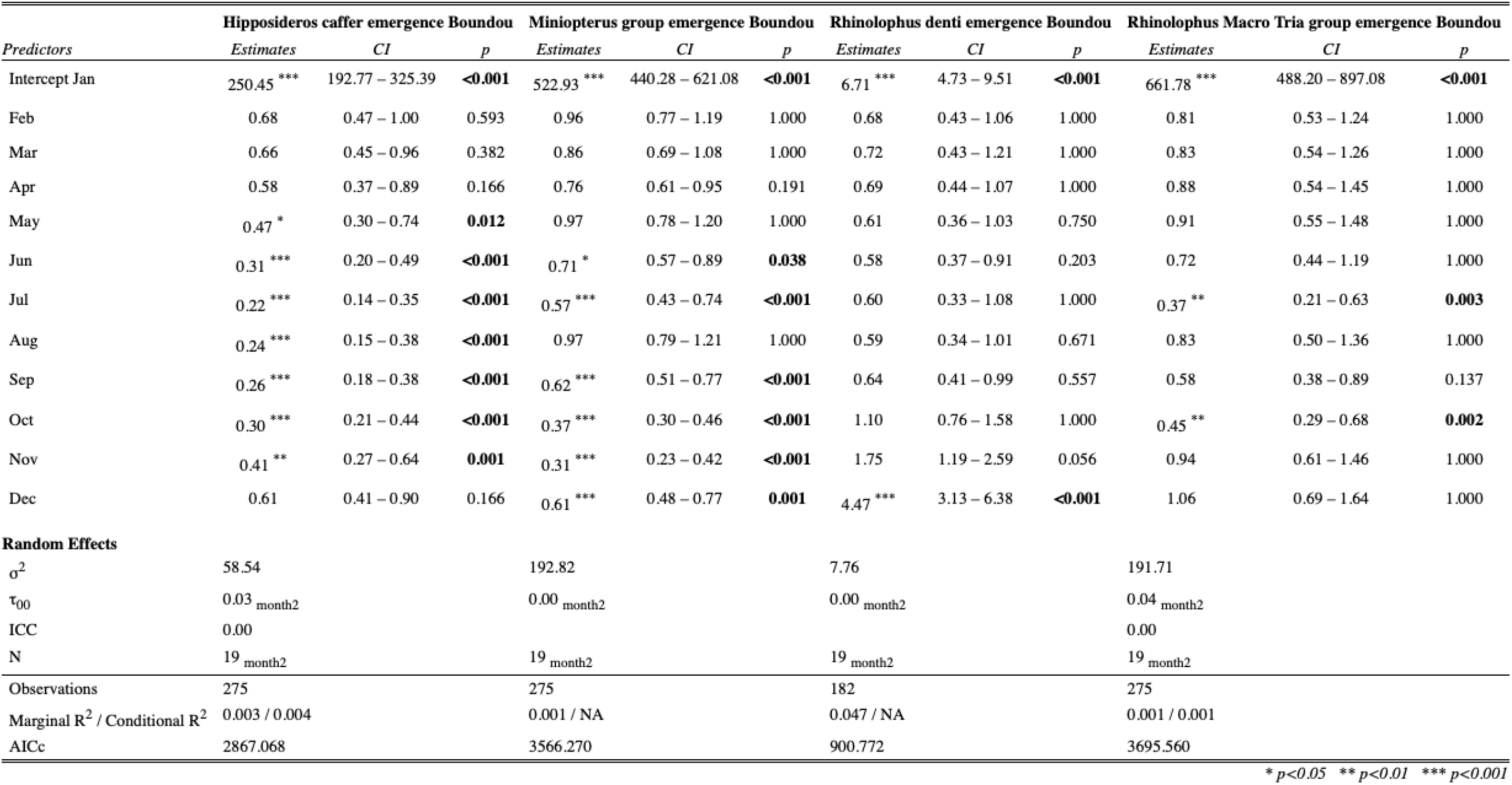
Generalized linear mixed (GLM) with Bonferroni correction describing the relationship between activity and month for the four acoustic groups in the Boundou cave during the emergence time of bats.

**Appendix 5.** Generalized linear mixed (GLM) with Bonferroni correction describing the relationship between activity and month for the four acoustic groups in the Mont Belo cave during the emergence time of bats.

| *Predictors* | **Hipposideros caffer emergence Mont Belo** |  |  | **Miniopterus group emergence Mont Belo** |  |  | **Rhino landeri-alcyone emergence Mont Belo** |  |  | **Rhinolophus denti emergence Mont Belo** |  |  |
| --- | --- | --- | --- | --- | --- | --- | --- | --- | --- | --- | --- | --- |
|  | *Estimates* | *CI* | *p* | *Estimates* | *CI* | *p* | *Estimates* | *CI* | *p* | *Estimates* | *CI* | *p* |
| Intercept Jan | 9.35 *** | 6.82 – 12.82 | <0.001 | 737.51 *** | 636.15 – 855.01 | <0.001 | 499.73 *** | 346.52 – 720.68 | <0.001 | 1.00 | NaN – NaN | NaN |
| Feb | 1.05 | 0.74 – 1.47 | 1.000 | 1.08 | 0.91 – 1.29 | 1.000 | 1.03 | 0.66 – 1.60 | 1.000 |  |  |  |
| Mar | 1.34 | 0.91 – 1.98 | 1.000 | 1.15 | 0.96 – 1.39 | 1.000 | 1.14 | 0.72 – 1.79 | 1.000 |  |  |  |
| Apr | 1.08 | 0.72 – 1.63 | 1.000 | 1.06 | 0.86 – 1.31 | 1.000 | 1.27 | 0.76 – 2.11 | 1.000 |  |  |  |
| May | 1.14 | 0.79 – 1.64 | 1.000 | 1.03 | 0.85 – 1.26 | 1.000 | 1.12 | 0.68 – 1.86 | 1.000 |  |  |  |
| Jun | 0.68 | 0.44 – 1.05 | 0.958 | 0.79 | 0.64 – 0.97 | 0.316 | 0.92 | 0.55 – 1.54 | 1.000 |  |  |  |
| Jul | 0.83 | 0.55 – 1.24 | 1.000 | 0.74 | 0.60 – 0.91 | 0.064 | 0.85 | 0.51 – 1.42 | 1.000 |  |  |  |
| Aug | 0.61 | 0.37 – 1.00 | 0.588 | 0.70 * | 0.56 – 0.86 | 0.012 | 0.77 | 0.46 – 1.28 | 1.000 |  |  |  |
| Sep | 0.76 | 0.52 – 1.11 | 1.000 | 0.73 * | 0.60 – 0.88 | 0.013 | 0.63 | 0.40 – 0.99 | 0.542 | 1.00 | NaN – NaN | NaN |
| Oct | 0.83 | 0.58 – 1.18 | 1.000 | 1.05 | 0.88 – 1.25 | 1.000 | 1.06 | 0.68 – 1.65 | 1.000 | 1.00 | NaN – NaN | NaN |
| Nov | 0.90 | 0.61 – 1.33 | 1.000 | 0.99 | 0.82 – 1.19 | 1.000 | 1.00 | 0.63 – 1.57 | 1.000 |  |  |  |
| Dec | 1.85 ** | 1.34 – 2.56 | 0.003 | 1.00 | 0.84 – 1.19 | 1.000 | 0.98 | 0.62 – 1.53 | 1.000 |  |  |  |
| **Random Effects** |  |  |  |  |  |  |  |  |  |  |  |  |
| $\sigma^2$ | 13.88 | | | 152.17 | | | 169.52 | | | 0.00 | | |
| $\tau_{00}$ | 0.00<sub>month2</sub> | | | 0.00<sub>month2</sub> | | | 0.03<sub>month2</sub> | | | 0.00<sub>month2</sub> | | |
| ICC |  |  |  | 0.00 |  |  | 0.00 |  |  |  |  |  |
| N | 18<sub>month2</sub> |  |  | 18<sub>month2</sub> |  |  | 18<sub>month2</sub> |  |  | 4<sub>month2</sub> |  |  |
| Observations | 271 |  |  | 293 |  |  | 293 |  |  | 4 |  |  |
| Marginal R<sup>2</sup> / Conditional R<sup>2</sup> | 0.007 / NA |  |  | 0.000 / 0.000 |  |  | 0.000 / 0.000 |  |  | 0.000 / NA |  |  |
| AICc | 1644.111 |  |  | 3796.908 |  |  | 3859.854 |  |  | NA |  |  |

\* $p < 0.05$ \*\* $p < 0.01$ \*\*\* $p < 0.001$

**Appendix 6.** Generalized linear mixed (GLM) with Bonferroni correction describing the relationship between activity and month for the four acoustic groups in the Boundou cave during the enter time of bats.

| *Predictors* | **Hipposideros caffer enter Boundou** |  |  | **Miniopterus group enter Boundou** |  |  | **Rhinolophus denti enter Boundou** |  |  | **Rhinolophus Macro Tria group enter Boundou** |  |  |
| --- | --- | --- | --- | --- | --- | --- | --- | --- | --- | --- | --- | --- |
|  | *Estimates* | *CI* | *p* | *Estimates* | *CI* | *p* | *Estimates* | *CI* | *p* | *Estimates* | *CI* | *p* |
| Intercept Jan | 249.41 *** | 156.62 – 397.20 | <0.001 | 463.67 *** | 314.74 – 683.07 | <0.001 | 17.20 *** | 11.53 – 25.65 | <0.001 | 439.89 *** | 289.90 – 667.50 | <0.001 |
| Feb | 1.07 | 0.59 – 1.96 | 1.000 | 1.71 | 1.07 – 2.74 | 0.290 | 0.43 * | 0.25 – 0.73 | 0.020 | 1.65 | 0.96 – 2.85 | 0.865 |
| Mar | 1.57 | 0.87 – 2.85 | 1.000 | 1.63 | 1.02 – 2.60 | 0.497 | 0.55 | 0.33 – 0.91 | 0.242 | 1.54 | 0.89 – 2.67 | 1.000 |
| Apr | 1.16 | 0.58 – 2.31 | 1.000 | 1.53 | 0.91 – 2.57 | 1.000 | 0.37 ** | 0.21 – 0.64 | 0.005 | 1.70 | 0.90 – 3.20 | 1.000 |
| May | 1.18 | 0.60 – 2.34 | 1.000 | 1.45 | 0.86 – 2.44 | 1.000 | 0.41 * | 0.24 – 0.72 | 0.021 | 1.86 | 0.99 – 3.50 | 0.656 |
| Jun | 0.75 | 0.38 – 1.49 | 1.000 | 0.76 | 0.44 – 1.31 | 1.000 | 0.29 *** | 0.16 – 0.50 | <0.001 | 1.26 | 0.66 – 2.40 | 1.000 |
| Jul | 0.48 | 0.24 – 0.96 | 0.457 | 0.89 | 0.52 – 1.52 | 1.000 | 0.31 *** | 0.17 – 0.55 | 0.001 | 1.50 | 0.80 – 2.84 | 1.000 |
| Aug | 0.46 | 0.22 – 0.93 | 0.377 | 1.36 | 0.81 – 2.30 | 1.000 | 0.36 ** | 0.20 – 0.65 | 0.007 | 1.40 | 0.74 – 2.65 | 1.000 |
| Sep | 0.49 | 0.27 – 0.89 | 0.229 | 1.17 | 0.74 – 1.87 | 1.000 | 0.29 *** | 0.17 – 0.50 | <0.001 | 1.02 | 0.59 – 1.76 | 1.000 |
| Oct | 0.67 | 0.37 – 1.21 | 1.000 | 1.19 | 0.75 – 1.89 | 1.000 | 0.48 * | 0.30 – 0.77 | 0.028 | 0.83 | 0.48 – 1.44 | 1.000 |
| Nov | 0.69 | 0.37 – 1.31 | 1.000 | 1.12 | 0.68 – 1.83 | 1.000 | 1.15 | 0.72 – 1.82 | 1.000 | 1.39 | 0.79 – 2.44 | 1.000 |
| Dec | 1.00 | 0.54 – 1.84 | 1.000 | 1.38 | 0.85 – 2.23 | 1.000 | 2.80 *** | 1.73 – 4.54 | <0.001 | 2.01 | 1.16 – 3.47 | 0.154 |
| **Random Effects** |  |  |  |  |  |  |  |  |  |  |  |  |
| $\sigma^2$ | 155.58 | | | 202.81 | | | 9.54 | | | 229.51 | | |
| $\tau_{00}$ | 0.06<sub>month2</sub> | | | 0.03<sub>month2</sub> | | | 0.02<sub>month2</sub> | | | 0.06<sub>month2</sub> | | |
| ICC | 0.00 |  |  | 0.00 |  |  | 0.00 |  |  | 0.00 |  |  |
| N | 19<sub>month2</sub> |  |  | 19<sub>month2</sub> |  |  | 19<sub>month2</sub> |  |  | 19<sub>month2</sub> |  |  |
| Observations | 276 |  |  | 277 |  |  | 242 |  |  | 277 |  |  |
| Marginal R<sup>2</sup> / Conditional R<sup>2</sup> | 0.001 / 0.001 |  |  | 0.000 / 0.000 |  |  | 0.042 / 0.044 |  |  | 0.000 / 0.001 |  |  |
| AICc | 3301.278 |  |  | 3769.686 |  |  | 1366.644 |  |  | 3790.904 |  |  |

\* $p < 0.05$ \*\* $p < 0.01$ \*\*\* $p < 0.001$

**Appendix 7.** Generalized linear mixed (GLM) with Bonferroni correction describing the relationship between activity and month for the four acoustic groups in the Mont Belo cave during the enter time of bats.

| *Predictors* | **Hipposideros caffer enter Mont Belo** |  |  | **Miniopterus group enter Mont Belo** |  |  | **Rhino landeri-alcyone enter Mont Belo** |  |  | **Rhinolophus denti enter Mont Belo** |  |  |
| --- | --- | --- | --- | --- | --- | --- | --- | --- | --- | --- | --- | --- |
|  | *Estimates* | *CI* | *p* | *Estimates* | *CI* | *p* | *Estimates* | *CI* | *p* | *Estimates* | *CI* | *p* |
| Intercept Jan | 20.98 *** | 12.75 – 34.52 | <0.001 | 570.35 *** | 501.54 – 648.60 | <0.001 | 663.65 *** | 530.64 – 830.00 | <0.001 | 1.17 | 0.47 – 2.93 | 1.000 |
| Feb | 1.30 | 0.72 – 2.33 | 1.000 | 1.22 | 1.05 – 1.41 | 0.086 | 1.05 | 0.80 – 1.38 | 1.000 |  |  |  |
| Mar | 1.41 | 0.77 – 2.58 | 1.000 | 1.17 | 0.99 – 1.38 | 0.822 | 0.92 | 0.69 – 1.23 | 1.000 |  |  |  |
| Apr | 1.08 | 0.54 – 2.15 | 1.000 | 1.27 | 1.07 – 1.50 | 0.063 | 1.19 | 0.87 – 1.62 | 1.000 |  |  |  |
| May | 0.83 | 0.42 – 1.63 | 1.000 | 1.37 *** | 1.18 – 1.59 | 0.001 | 1.09 | 0.80 – 1.49 | 1.000 | 1.00 | 0.32 – 3.13 | 1.000 |
| Jun | 0.54 | 0.27 – 1.08 | 0.988 | 1.22 | 1.04 – 1.43 | 0.153 | 0.87 | 0.64 – 1.19 | 1.000 |  |  |  |
| Jul | 0.50 | 0.24 – 1.03 | 0.705 | 1.21 | 1.03 – 1.42 | 0.229 | 0.66 | 0.48 – 0.92 | 0.155 |  |  |  |
| Aug | 0.38 | 0.18 – 0.78 | 0.103 | 0.94 | 0.79 – 1.11 | 1.000 | 0.52 *** | 0.37 – 0.72 | 0.001 |  |  |  |
| Sep | 0.52 | 0.28 – 0.96 | 0.453 | 0.95 | 0.81 – 1.11 | 1.000 | 0.61 ** | 0.46 – 0.81 | 0.007 |  |  |  |
| Oct | 0.66 | 0.36 – 1.20 | 1.000 | 0.94 | 0.81 – 1.10 | 1.000 | 0.73 | 0.55 – 0.96 | 0.279 |  |  |  |
| Nov | 0.64 | 0.34 – 1.20 | 1.000 | 0.79 | 0.66 – 0.94 | 0.110 | 0.63 * | 0.47 – 0.84 | 0.021 |  |  |  |
| Dec | 1.22 | 0.66 – 2.25 | 1.000 | 0.98 | 0.84 – 1.14 | 1.000 | 0.97 | 0.73 – 1.29 | 1.000 | 1.55 | 0.53 – 4.57 | 1.000 |
| **Random Effects** |  |  |  |  |  |  |  |  |  |  |  |  |
| $\sigma^2$ | 22.89 | | | 166.33 | | | 157.22 | | | | | |
| $\tau_{00}$ | 0.04<sub>month2</sub> | | | 0.00<sub>month2</sub> | | | 0.01<sub>month2</sub> | | | | | |
| ICC | 0.00 |  |  |  |  |  | 0.00 |  |  |  |  |  |
| N | 18<sub>month2</sub> |  |  | 18<sub>month2</sub> |  |  | 18<sub>month2</sub> |  |  | 4<sub>month2</sub> |  |  |
| Observations | 279 |  |  | 294 |  |  | 294 |  |  | 4 |  |  |
| Marginal R<sup>2</sup> / Conditional R<sup>2</sup> | 0.007 / 0.009 |  |  | 0.000 / NA |  |  | 0.000 / 0.000 |  |  | NA |  |  |
| AICc | 1995.258 |  |  | 3840.501 |  |  | 3818.869 |  |  | -12.700 |  |  |

\* $p < 0.05$ \*\* $p < 0.01$ \*\*\* $p < 0.001$

**Appendix 8.**
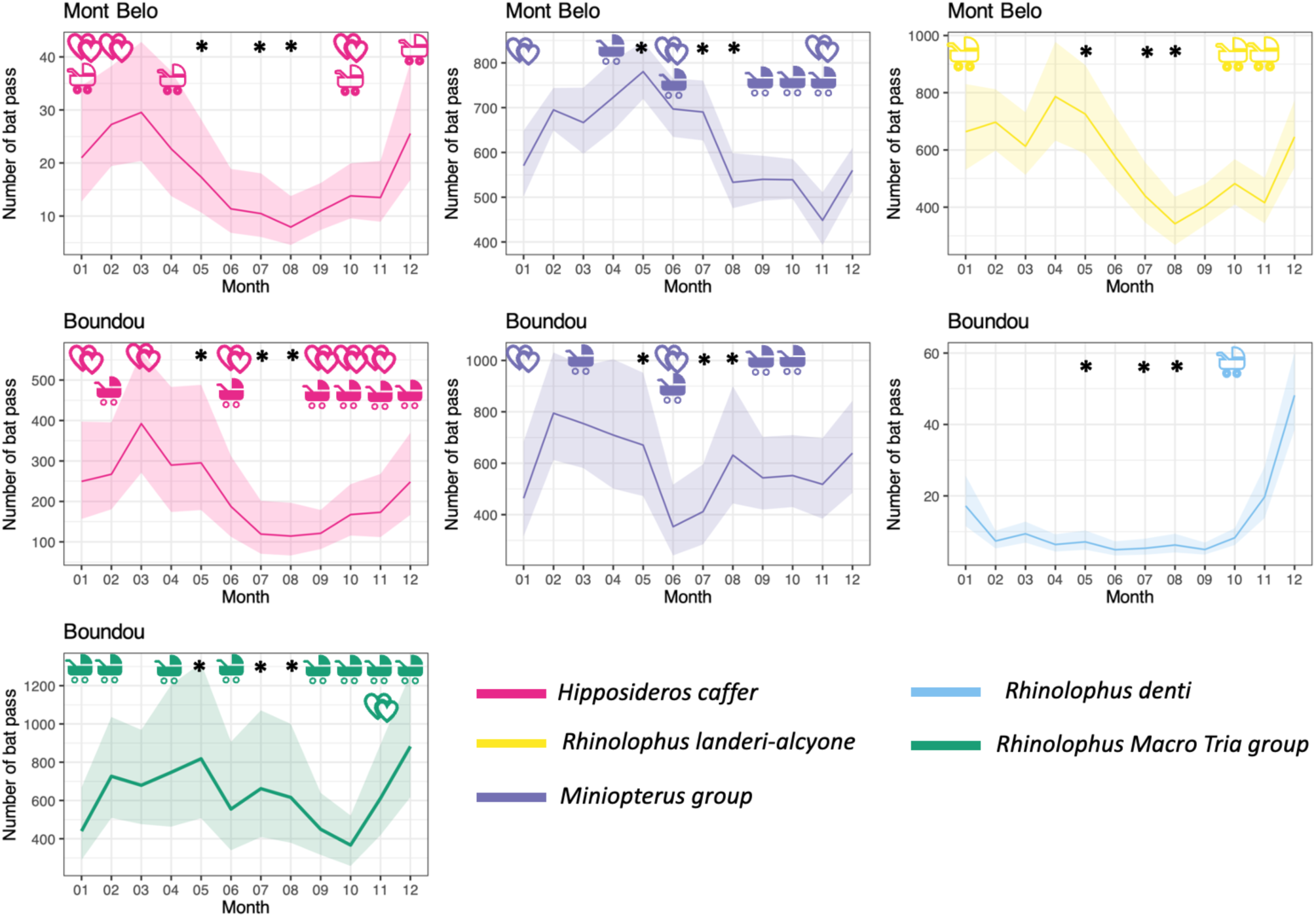
Prediction of the activity of acoustic group entries with 95% confidence interval as a function of months and study sites (Boundou and Mont Belo cave). The Rhinolophus denti group at Mont Belo has not been represented here due to lack of data. The reproductive status of the species or acoustic group is indicated by two pictograms with the color of the acoustics groups: 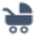 female reproductive period (lactation, pregnant, young rearing) and 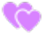: mating period/male activity. The months of May, July and August, which have an *, are month when no captures were made, hence the absence of data on the reproductive status of the acoustic group. Outlined icons indicate a suggested (S) or insufficient (I) confidence level; filled icons indicate confirmed (C).

## Acknowledgements

We would also like to extend our warmest thanks to Léa Mariton for her support and advice on the R scripts for the density of bat activity over time. Many thanks to our Congolese partners: the “Laboratoire National de Santé Publique de Brazzaville” (LNSP), the “Direction Générale de l’Elevage” (DGE) and “Ministère de l’économie forestière” for their support in this project, their help with the fieldwork and the applications for the capture and ethics permits. Thanks also to the bat field team who helped collect data: F. Nguilili, C. Bazola, N. Nguimbi (our wizard), R. Dimoukissi and R. Nguimbi. Finally, we would like to thank the local authorities in Dolisie and the local population for their help with this project.

## Data, scripts, code, and supplementary information availability

The summary file of species and the scripts used to create the figures can be obtained by following this link: https://doi.org/10.5281/zenodo.15636144

## References

Alcock, J. (2013). Animal Behavior: An Evolutionary Approach (10th edition). Sinauer Associates.

Appel, G., da Rocha, P.A., Cerqueira, P.V. et al. (2025). Using active and passive methods to monitor cave-dwelling bats in Amazonian caves. Biodiversity Conservation, 34, 4429–4449 10.1007/s10531-025-03160-4

Appel, G., Capaverde, U. D., De Oliveira, L. Q., do Amaral Pereira, L. G., da Cunha Tavares, V., López-Baucells, A., Magnusson, W. E., Baccaro, F. B., & Bobrowiec, P. E. (2021). Use of complementary methods to sample bats in the Amazon. Acta Chiropterologica, 23(2), 499–511. 10.3161/15081109ACC2021.23.2.017

Appel, G., López-Baucells, A., Magnusson, W. E., & Bobrowiec, P. E. D. (2019). Temperature, rainfall, and moonlight intensity effects on activity of tropical insectivorous bats. Journal of Mammalogy, 100(6), 1889–1900. 10.1093/jmammal/gyz140

Azmy, S. N., Sah, S. A. M., Shafie, N. J., Ariffin, A., Majid, Z., Ismail, M. N. A., & Shamsir, M. S. (2012). Counting in the dark: Non-intrusive laser scanning for population counting and identifying roosting bats. Scientific Reports, 2(1), 1–4.10.1038/srep00524

Balogh, A., Ngo, L., Zigler, K. S., & Dixon, G. (2020). Population genomics in two cave-obligate invertebrates confirms extremely limited dispersal between caves. Scientific Reports, 10(1), 17554.10.1038/s41598-020-74508-9

Barclay, R. M., & Kurta, A. (2007). Ecology and behavior of bats roosting in tree cavities and under bark. Bats in Forests: Conservation and Management (MJ Lacki, JP Hayes, and A. Kurta Eds.). Johns Hopkins University Press, Baltimore, Maryland, 352, 17–59.

Barnhart, P. R., & Gillam, E. H. (2014). The impact of sampling method on maximum entropy species distribution modeling for bats. Acta Chiropterologica, 16(1), 241–248.10.3161/150811014X683435

Barnosky, A. D., Matzke, N., Tomiya, S., Wogan, G. O. U., Swartz, B., Quental, T. B., Marshall, C., McGuire, J. L., Lindsey, E. L., Maguire, K. C., Mersey, B., & Ferrer, E. A. (2011). Has the Earth’s sixth mass extinction already arrived? Nature, 471(7336), 51–57. 10.1038/nature09678

Barros, J. D. S., Bernard, E., & Ferreira, R. L. (2020). Ecological preferences of neotropical cave bats in roost site selection and their implications for conservation. Basic and Applied Ecology, 45, 31–41. 10.1016/j.baae.2020.03.007

Bates, P. J. J., Cameron, K., Pearch, M. J., & Hayes, B. (2013). A review of the bats (Chiroptera) of the republic of Congo, including eight species new to the country. Acta Chiropterologica, 15(2), 313–340. 10.3161/150811013X678955

Bernard, R. & Meester, J. A. J. (1982). Female reproduction and the female reproductive cycle of Hipposideros caffer caffer (Sundevall, 1846) in Natal, South Africa. Annals of the Transvaal Museum, 33(8), 131–144.

Bion, R. (2023). ggradar: Create radar charts using ggplot2. R Package Version 0.2.

Brooks, M. E., Kristensen, K., Van Benthem, K. J., Magnusson, A., Berg, C. W., Nielsen, A., Skaug, H. J., Machler, M., & Bolker, B. M. (2017). glmmTMB balances speed and flexibility among packages for zero-inflated generalized linear mixed modeling. The R Journal, 9(2), 378–400. 10.32614/RJ-2017-066

Brosset, A. (1968). La Permutation du Cycle sexuel saisonnier chez le chiroptère Hipposideros caffer, au voisinage de l’Équateur.

Browning, E., Gibb, R., Glover-Kapfer, P., & Jones, K. E. (2017). Passive acoustic monitoring in ecology and conservation. 10.25607/OBP-876

Bullock, D., Combes, B., Eales, L., & Pritchard, J. (1987). Analysis of the timing and pattern of emergence of the pipistrelle bat (Pipistrellus pipistrettus). Journal of Zoology, 211(2), 267–274. 10.1111/j.1469-7998.1987.tb01533.x

Cabin, R. J., & Mitchell, R. (2000). To Bonferroni or not to Bonferroni: When and how are the questions. ESA Bulletin, 81, 246–248. 10.2307/20168454

Catto, C., Racey, P., & Stephenson, P. (1995). Activity patterns of the serotine bat (Eptesicus serotinus) at a roost in southern England. Journal of Zoology, 235(4), 635–644. 10.1111/j.1469-7998.1995.tb01774.x

Ceballos, G., Ehrlich, P. R., Barnosky, A. D., García, A., Pringle, R. M., & Palmer, T. M. (2015). Accelerated modern human–induced species losses: Entering the sixth mass extinction. Science Advances, 1(5), e1400253. 10.1126/sciadv.1400253

Cumming, G., & Bernard, R. (1997). Rainfall, food abundance and timing of parturition in African bats. Oecologia, 111, 309–317. 10.1007/s004420050240

Deleva, S., Toshkova, N., Kolev, M., & Tanalgo, K. C. (2023). Important underground roosts for bats in Bulgaria: current state and priorities for conservation. Biodiversity Data Journal, 11, e98734.

Dietz, M., & Kalko, E. K. (2006). Seasonal changes in daily torpor patterns of free-ranging female and male Daubenton’s bats (Myotis daubentonii). Journal of Comparative Physiology B, 176, 223–231. 10.1007/s00360-005-0043-x

Erkert, H. G. (1982). Ecological Aspects of Bat Activity Rhythms. In T. H. Kunz (Ed.), Ecology of Bats (pp. 201–242). Springer US. 10.1007/978-1-4613-3421-7_5

Flaquer, C., Torre, I., & Arrizabalaga, A. (2007). Comparison of sampling methods for inventory of bat communities. Journal of Mammalogy, 88(2), 526–533. 10.1644/06-MAMM-A-135R1.1

Forsyth, D. M., Scroggie, M. P., & McDonald-Madden, E. (2006). Accuracy and precision of grey-headed flying-fox (Pteropus poliocephalus) flyout counts. Wildlife Research, 33(1), 57–65. 10.1071/WR05029

Furey, N. M., & Racey, P. A. (2016). Conservation ecology of cave bats. In Bats in the Anthropocene: Conservation of Bats in a Changing World (pp. 463–500). Springer, Cham. 10.1007/978-3-319-25220-9_15

Gibb, R., Browning, E., Glover-Kapfer, P., & Jones, K. E. (2019). Emerging opportunities and challenges for passive acoustics in ecological assessment and monitoring. Methods in Ecology and Evolution, 10(2), 169–185. 10.1111/2041-210X.13101

Greville, L. J., Bueno, L. M., Pollock, T., & Faure, P. A. (2022). Quantification of Urinary Sex Steroids in the Big Brown Bat (Eptesicus fuscus). Physiological and Biochemical Zoology, 95(1), 22–34.

Günther, L., Lopez, M. D., Knörnschild, M., Reid, K., Nagy, M., & Mayer, F. (2016). From resource to female defence: The impact of roosting ecology on a bat’s mating strategy. Royal Society Open Science, 3(11), 160503. 10.1098/rsos.160503

Happold, D.C.D. and Happold, M. (1990), Reproductive strategies of bats in Africa. Journal of Zoology, 222: 557–583. 10.1111/j.1469-7998.1990.tb06014.x

Hartig, F., & Hartig, M. (2022). Package ‘DHARMa’. R Package. Available Online: Https://CRAN. R-Project. Org/Package= DHARMa (Accessed on 5 September 2022).

Henry, M., Thomas, D. W., Vaudry, R., & Carrier, M. (2002). Foraging Distances and Home Range of Pregnant and Lactating Little Brown Bats (Myotis Lucifugus). Journal of Mammalogy, 83(3), 767–774. 10.1644/1545-1542(2002)083<0767:FDAHRO>2.0.CO;2

Hill, A. P., Prince, P., Snaddon, J. L., Doncaster, C. P., & Rogers, A. (2019). AudioMoth: A low-cost acoustic device for monitoring biodiversity and the environment. HardwareX, 6, e00073. 10.1016/j.ohx.2019.e00073

Hughes, A. C. (2017). Understanding the drivers of Southeast Asian biodiversity loss. Ecosphere, 8(1), e01624. 10.1002/ecs2.1624

IPBES (2019): Global assessment report on biodiversity and ecosystem services of the Intergovernmental Science-Policy Platform on Biodiversity and Ecosystem Services. E. S. Brondizio, J. Settele, S. Díaz, and H. T. Ngo (editors). IPBES secretariat, Bonn, Germany. 1148 pages. 10.5281/zenodo.3831673

IUCN, N. and I. (International U. for C. of N. (2023). The IUCN Red List of Threatened Species. Version 2022-2. IUCN Red List of Threatened Species. https://www.iucnredlist.org/en

Jenber, A. J., & Wili, G. G. (2021). Insect diversity in wet and dry seasons at hursa forest, central Ethiopia. International Journal of Entomology Research, 6(4), 59–63.

Jones, G., & Rydell, J. (1994). Foraging strategy and predation risk as factors influencing emergence time in echolocating bats. Philosophical Transactions of the Royal Society of London. Series B: Biological Sciences, 346(1318), 445–455. 10.1098/rstb.1994.0161

Keeley, B. W., & Tuttle, M. D. (1999). Bats in American bridges. Bat Conservation International Incorporated.

Koger, B., Hurme, E., Costelloe, B. R., O’Mara, M. T., Wikelski, M., Kays, R., & Dechmann, D. K. N. (2023). An automated approach for counting groups of flying animals applied to one of the world’s largest bat colonies. Ecosphere, 14(6), e4590. 10.1002/ecs2.4590

Kumar, V., & Umapathy, G. (2019). Non-invasive monitoring of steroid hormones in wildlife for conservation and management of endangered species-A review. Indian J Exp Biol, 57, 307–314.

Kunz, T. H. (1982). Roosting ecology of bats. In Ecology of bats (pp. 1–55). Springer.

Kunz, T. H., Lumsden, L. F., & Fenton, M. (2003). Ecology of cavity and foliage roosting bats. Bat Ecology, 1, 3–89. 10.5281

Kurta, A., Bell, G. P., Nagy, K. A., & Kunz, T. H. (1989). Energetics of pregnancy and lactation in freeranging little brown bats (Myotis lucifugus). Physiological Zoology, 62(3), 804–818. 10.1111/j.1469-7998.1992.tb04433.x

Labadie, M., Morand, S., Bourgarel, M., Niama, F. R., Nguilili, F., N’Kaya, T., Caron, A., De Nys, H., (2025a). PeerJ. Habitat sharing and interspecies interactions in caves used by bats in the Republic of Congo 13:e18145 10.7717/peerj.18145

Labadie, M., Morand, S., Caron, A., De Nys, H. M., Niama, F. R., Nguilili, F.,… Roemer, C. (2025b). Passive acoustic monitoring of cave-dwelling bats with a sonotype classifier. Bioacoustics, 1–29. 10.1080/09524622.2024.2438600

Labadie, M. (2025c). Complete results of acoustic data collected in the Republic of the Congo (Tadarida classifier). Zenodo. 10.5281/zenodo.15636144

Lewis, S. E. (1995). Roost Fidelity of Bats: A Review. Journal of Mammalogy, 76(2), 481–496. 10.2307/1382357

Maier, C. (1992). Activity patterns of pipistrelle bats (Pipistrellus pipistrellus) in Oxfordshire. Journal of Zoology, 228(1), 69–80.

Mariton, L., Le Viol, I., Bas, Y., & Kerbiriou, C. (2023). Characterising diel activity patterns to design conservation measures: Case study of European bat species. Biological Conservation, 277, 109852. 10.1016/j.biocon.2022.109852

Mcwilliam, A.N. (1988), The reproductive cycle of male long-fingered bats, Miniopterus minor (Chiroptera: Vespertilionidae), in a seasonal environment of the African Tropics. Journal of Zoology, 216: 119–129. 10.1111/j.1469-7998.1988.tb02419.x

Medellin, R. A., Wiederholt, R., & Lopez-Hoffman, L. (2017). Conservation relevance of bat caves for biodiversity and ecosystem services. Biological Conservation, 211, 45–50.

Meierhofer, M. B., Johnson, J. S., Perez-Jimenez, J., Ito, F., Webela, P. W., Wiantoro, S., Bernard, E., Tanalgo, K. C., Hughes, A., Cardoso, P., Lilley, T., & Mammola, S. (2024). Effective conservation of subterranean-roosting bats. Conservation Biology, 38(1), e14157. 10.1111/cobi.14157

Mering, E. D., & Chambers, C. L. (2014). Thinking outside the box: A review of artificial roosts for bats. Wildlife Society Bulletin, 38(4), 741–751. 10.1002/wsb.461

Mickleburgh, S. P., Hutson, A. M., & Racey, P. A. (2002). A review of the global conservation status of bats. Oryx, 36(1), 18–34. doi:10.1017/S0030605302000054

Millon, L., Julien, J.-F., Julliard, R., & Kerbiriou, C. (2015). Bat activity in intensively farmed landscapes with wind turbines and offset measures. Ecological Engineering, 75, 250–257. 10.1016/j.ecoleng.2014.11.050

Monadjem, A., Taylor, P. J., & Schoeman, M. C. (2020). Bats of southern and central Africa: A biogeographic and taxonomic synthesis. Wits University Press.

Moutaouakil, S., Souza-Silva, M., Oliveira, L. F., Ghamizi, M., & Ferreira, R. L. (2024). A cave with remarkably high subterranean diversity in Africa and its significance for biodiversity conservation. Subterranean Biology, 50, 1–28.

Nkoana, T. T. (2020). Temporal changes in food resource availability between two karst bat assemblages (Doctoral dissertation, University of Pretoria).

Neuhuber, S., Plan, L., Gier, S., Hintersberger, E., Lachner, J., Scholz, D., Lüthgens, C., Braumann, S., Bodenlenz, F., & Voit, K. (2020). Numerical age dating of cave sediments to quantify vertical movement at the Alpine-Carpathian transition in the Plio-and Pleistocene. Geologica Carpathica, 71(6), 539–557. 10.31577/GeolCarp.71.6.5

Okonkwo, E. E., Afoma, E., & Martha, I. W. (2017). Cave Tourism and its Implications to Tourism Development in Nigeria: A Case Study of Agu-Owuru Cave in Ezeagu. International Journal of Research, 3, 16–24. doi:10.20431/2455-0043.0303003

Ormsbee, P. C., Kiser, J. D., & Perlmeter, S. I. (2007). Importance of night roosts to the ecology of bats. Bats in Forests: Conservation and Management. Johns Hopkins University Press, Baltimore, MD, 129–151.

Patriquin, K. J., & Ratcliffe, J. M. (2016). Should I stay or should I go? Fission–fusion dynamics in bats. Sociality in Bats, 65–103. 10.1007/978-3-319-38953-0_4

Patterson, B. D., Dick, C. W., & Dittmar, K. (2007). Roosting habits of bats affect their parasitism by bat flies (Diptera: Streblidae). Journal of Tropical Ecology, 23(2), 177–189. doi:10.1017/S0266467406003816

Pretorius, M., Markotter, W. & Keith, M. Assessing the extent of land-use change around important bat-inhabited caves. BMC Zool 6, 31 (2021). 10.1186/s40850-021-00095-5

Postawa, T., & Gas, A. (2009). Do the thermal conditions in maternity colony roost determine the size of young bats? Comparison of attic and cave colonies of Myotis myotis in Southern Poland. Folia Zoologica, 58(4), 396.

R Core Team. (2023). R: A Language and Environment for Statistical Computing. R Foundation for Statistical Computing. https://www.R-project.org/

Racey, P. A., & Entwistle, A. C. (2000). Life-history and Reproductive Strategies of Bats. In Reproductive Biology of Bats (pp. 363–414). Elsevier. 10.1016/B978-012195670-7/50010-2

Revilla-Martín, N., Budinski, I., Puig-Montserrat, X., Flaquer, C., & López-Baucells, A. (2020). Monitoring cave-dwelling bats using remote passive acoustic detectors: A new approach for cave monitoring. Bioacoustics, 00(00), 1–16. 10.1080/09524622.2020.1816492

Roemer, C., Julien, J. F., & Bas, Y. (2021). An automatic classifier of bat sonotypes around the world. Methods in Ecology and Evolution, 12(12), 2432–2444. 10.1111/2041-210X.13721

Russo, D., & Ancillotto, L. (2015). Sensitivity of bats to urbanization: A review. Mammalian Biology, 80(3), 205–212. 10.1016/j.mambio.2014.10.003

Racey, P.A. & Entwistle, A.C. (2000). Conservation Ecology of Cave Bats. In: Conservation of Tropical Biodiversity. Springer.

Rydell, J. (1993). Variation in Foraging Activity of an Aerial Insectivorous Bat during Reproduction. Journal of Mammalogy, 74(2), 503–509. 10.2307/1382411

Sabol, B. M., & Hudson, M. K. (1995). Technique using thermal infrared-imaging for estimating populations of gray bats. Journal of Mammalogy, 76(4), 1242–1248. 10.2307/1382618

Simmons, N. B., & Cirranello, A. L. (2025). Bat Species of the World: A taxonomic and geographic database. https://batnames.org/query.html

Simons, J. W. (1998). Guano mining in Kenyan lava tunnel caves. International Journal of Speleology, 27, 4. https://api.semanticscholar.org/CorpusID:56146083

Struebig, M. J., Kingston, T., Zubaid, A., Le Comber, S. C., Mohd-Adnan, A., Turner, A., Kelly, J., Bożek, M., & Rossiter, S. J. (2009). Conservation importance of limestone karst outcrops for Palaeotropical bats in a fragmented landscape. Biological Conservation, 142(10), 2089–2096. 10.1016/j.biocon.2009.04.005

Sugai, L. S. M., Desjonquères, C., Silva, T. S. F., & Llusia, D. (2020). A roadmap for survey designs in terrestrial acoustic monitoring. Remote Sensing in Ecology and Conservation, 6(3), 220–235. 10.1002/rse2.131

Swift, S. M. (1980). Activity patterns of pipistrelle bats (Pipistrellus pipistrellus) in north-east Scotland. Journal of Zoology, 190(3), 285–295. 10.1111/j.1469-7998.1980.tb01428.x

Swift, S. M. (1997). Roosting and foraging behaviour of Natterer’s bats (Myotis nattereri) close to the northern border of their distribution. Journal of Zoology, 242(2), 375–384. 10.1111/j.1469-7998.1997.tb05809.x

Tanalgo, K. C., Oliveira, H. F., & Hughes, A. C. (2022). Mapping global conservation priorities and habitat vulnerabilities for cave-dwelling bats in a changing world. Science of The Total Environment, 843, 156909.

Tanalgo, K. C., & Hughes, A. C. (2024). Bat Cave Vulnerability Index 3.0 (BCVI-S): An integrative and scalable tool to prioritise bat caves for conservation. Global Ecology and Conservation, 57, e03396. 10.1016/j.gecco.2024.e03396

The Protection of Bat Roost Guidelines Subcommittee, Sheffield, S. R., Shaw, J. H., Heidt, G. A., & McClenaghan, L. R. (1992). Guidelines for the protection of bat roosts. Journal of Mammalogy, 707–710. 10.2307/1382051

Thieurmel, B., Elmarhraoui, A., & Thieurmel, M. B. (2022). Package ‘suncalc’. R Package Version 0.5. https://cran.r-project.org/package=suncalc

Thomas, D. W., & West, S. D. (1989). Sampling methods for bats. U.S. Department of Agriculture, Forest Service, Pacific Northwest Research Station. 10.2737/pnw-gtr-243

Voigt, C. C., & Schwarzenberger, F. (2008). Reproductive endocrinology of a small tropical bat (female Saccopteryx bilineata; Emballonuridae) monitored by fecal hormone metabolites. Journal of Mammalogy, 89(1), 50–57.

Westcott, D. A., & McKeown, A. (2004). Observer error in exit counts of flying-foxes (Pteropus spp.). Wildlife Research, 31(5), 551–558. 10.1071/WR03091

Whiting, J. C., Doering, B., Aho, K., & Rich, J. (2021). Long-term patterns of cave-exiting activity of hibernating bats in western North America. Scientific Reports, 11(1), 8175.

Wickham, H. (2016). ggplot2: Elegant Graphics for Data Analysis. Springer-Verlag New York. https://ggplot2.tidyverse.org

Wilkinson, G. S., Adams, D. M., & Rayner, J. G. (2024). Sex, season, age and status influence urinary steroid hormone profiles in an extremely polygynous neotropical bat. Hormones and Behavior, 164, 105606.

Wolda, H. (1988). Insect seasonality: Why? Annual Review of Ecology and Systematics, 19(1), 1–18. 10.1146/annurev.es.19.110188.000245

Wolkovich, E. M., Cook, B. I., McLauchlan, K. K., & Davies, T. J. (2014). Temporal ecology in the Anthropocene. Ecology Letters, 17(11), 1365–1379. 10.1111/ele.12353

Wright, G. S. (2009). Hipposideros caffer (Chiroptera: Hipposideridae). Mammalian Species, 845, 1–9. 10.1644/845.1.Key

